# Antibody treatment targeting nitrated alpha-synuclein counteracts protein spreading pathology

**DOI:** 10.64898/2026.05.21.726933

**Authors:** Ayse Ulusoy, Sarah Wright, Pietro La Vitola, Katharina Klinger, Eugenia Harbachova, Angela Rollar, Xiang Xu, Anushka Takhi, Nathan Behrendt, Anthony Mastracci, Bryan Lewis, Valerie Chen, Harry Ischiropoulos, Sheerin Shahidi-Latham, Irene Griswold-Prenner, Donato A. Di Monte

## Abstract

α-Synuclein nitration is a prominent post-translational modification in Parkinson’s disease, but whether nitrated α-synuclein merely reflects oxidative stress or actively contributes to pathology remains unclear. Here, we generated and characterized 6G6, an antibody selective for Tyr39-nitrated α-synuclein, and tested whether targeting this modified α-synuclein species affected pathology in different mouse models of α-synuclein aggregation and spread. In two models of α-synuclein overexpression targeting medullary vagal neurons, oxidative stress was induced by either exposure to the herbicide paraquat or transgenic heterozygous expression of the *Gba1*-L444P mutation. Both conditions were characterized by robust α-synuclein spreading that was markedly counteracted by 6G6 administration. A third model consisted of an injection of α-synuclein fibrils into the striatum of α-synuclein-overexpressing mice. In this model, treatment with 6G6 protected against fibril-induced aggregate pathology and ensuing degeneration of nigral dopaminergic neurons. In a pilot human study, CSF levels of Tyr39-nitrated α-synuclein were measured and found increased in Parkinson patients as compared to controls. These findings identify Tyr39-nitrated α-synuclein as a pathogenic, therapeutically targetable α-synuclein species linking oxidative/nitrative stress to PD pathological processes.

## INTRODUCTION

Parkinson’s disease (PD) is characterized by an abnormal accumulation of aggregated α-synuclein (αSyn) that forms intraneuronal inclusions called Lewy bodies and Lewy neurites.^1–3^ The brain distribution of Lewy inclusions follows a stereotypical pattern of progressive diffusion that affects anatomically interconnected brain regions. This spatiotemporal feature initially suggested that pathological forms of αSyn may spread from neuron to neuron and brain region to brain region, thus contributing to disease pathogenesis and advancement.^3–6^ Subsequent studies provided experimental evidence in support of this concept. In the *in vivo* setting, αSyn interneuronal transfer and brain spreading have been reproduced and investigated using primarily two rodent models. The first paradigm involves an intraparenchymal brain injection of pre-formed αSyn fibrils (PFFs) generated from the assembly of monomeric rodent or human αSyn (PFF model).^7,8^ Pathological αSyn aggregates are detected using an antibody against phosphorylated αSyn (P-αSyn) and, once formed in proximity of the injection site, progressively advance targeting interconnected brain regions. PFF-induced αSyn pathology has been described in both wild-type animals and transgenic mice overexpressing αSyn, with the latter typically displaying more severe and extensive P-αSyn lesions.^9–12^ The second *in vivo* model consists of an injection of adeno-associated viral vectors (AAVs) encoding human αSyn (HαSyn) into the rodent vagus nerve (AAV vagal model).^13,14^ This treatment induces HαSyn overexpression in the dorsal medulla oblongata (dMO), in particular the dorsal motor nucleus of the vagus nerve (DMnX), site of origin of vagal efferents, and the nucleus of the tractus solitarius (NTS), where vagal afferents terminate. Based on anatomical considerations, AAV-induced accumulation of the exogenous human protein should remain confined within these regions.^15,16^ Numerous studies using this paradigm have instead revealed a progressive caudo-rostral spreading of HαSyn that involved its transfer from “donor” neurons in the dMO to “recipient” axons innervating this brain region.^16,17^ The likely role of αSyn spreading in PD development bears important pathogenetic and therapeutic implications. It also raises key, yet unanswered questions. In particular, whether specific molecular mechanisms and distinct αSyn species are capable of prompting or modulating interneuronal protein transfer is poorly understood. Similarly, feasibility and effectiveness of therapeutic strategies targeting αSyn species with greater interneuronal mobility remain virtually untested.

Oxidative/nitrative stress has long been thought to play a role in PD pathogenetic processes and, in PD brain, clear demonstration of intraneuronal nitrative reactions is provided by the accumulation of nitrated αSyn within Lewy inclusions.^18–21^ Nitration may occur at one or more of the 4 αSyn’s tyrosine residues although, under pro-oxidant conditions, the Tyr39 site may be preferentially targeted.^22^ Nitration alters the structural and biophysical properties of αSyn, affects its interaction with membranes and has been associated with increased aggregation propensity and cytotoxicity.^23–25^ More recently, experimental work has indicated an intriguing relationship between neuronal oxidative stress, formation and accumulation of nitrated αSyn and enhanced neuron-to-neuron αSyn transfer. Using the AAV vagal model, such a relationship was consistently found in mice exposed to the pro-oxidant herbicide paraquat, mice with chemogenetically induced neuronal hyperactivity, and transgenic mice overexpressing αSyn on the background of a heterozygous mutation of the *Gba1* gene.^17,26,27^ Results of *in vitro* experiments using bimolecular fluorescence complementation to assess cell-to-cell αSyn transfer have also supported a direct role of nitrated αSyn in mediating intercellular αSyn exchanges during oxidative stress conditions.^26^ αSyn nitration can not only occur as a result of nonspecific chemical reactions but also involve a selective enzymatic mechanism catalyzed by glyoxalase domain-containing protein 4 (GLOD4).^28^ Interestingly, GLOD4 deficiency has been reported to significantly reduce the production of nitrated αSyn and, at the same time, counteract αSyn aggregation and the spreading of αSyn aggregate pathology in both *in vitro* and *in vivo* experimental models.^28,29^

Based on the above evidence supporting a harmful role of protein nitration, specific experiments were designed here to test the hypothesis that targeting this nitration, particularly at Tyr39, would prevent αSyn pathological spreading. To achieve this goal, we first developed and characterized a novel antibody, named 6G6, that selectively recognizes nitrated αSyn and displays high positional specificity for Tyr39-nitrated αSyn. The effects of systemic 6G6 administration were then assessed in mice with AAV- or PFF-induced spreading pathology.

Data revealed that, in the AAV vagal model, caudo-rostral HαSyn spreading was markedly reduced after 6G6 antibody treatment. 6G6 was also very effective in counteracting the propagation of αSyn pathology when PFFs were injected into the striatum of transgenic mice overexpressing HαSyn; in this latter model, the reduction of P-αSyn lesions by 6G6 administration also resulted in a significant decrease in neuronal cell damage. A final set of analyses was carried out to support the translational relevance of investigations focusing on nitrated αSyn as an important player in PD pathogenesis and a promising target for PD therapeutics. Tyr39-nitrated αSyn was measured in human CSF samples and, consistent with its generation and potential involvement in deleterious diseases processes, its levels were found to be markedly enhanced in PD patients as compared to healthy controls.

## Results

### Antibody development and characterization

To selectively target Tyr39-nitrated αSyn, a panel of antibodies binding this post-translationally modified protein was generated and screened. Several candidate clones were identified, including 6G6 and 8C2. In an initial peptide screen, these two antibodies were tested against a nitrated nine-mer peptide spanning residues 35-43 of αSyn and the corresponding non-nitrated peptide. Both 6G6 and 8C2 bound specifically to the nitrated nine-mer and did not recognize the non-nitrated peptide, consistent with their ability to discriminate nitrated from unmodified Tyr39. Clone 8C2 was further developed and used as an ELISA reagent (Fig. S1), while clone 6G6 underwent detailed characterization and was ultimately selected for anti-Tyr39-nitrated αSyn treatment of experimental animals. Initial tests were carried out to determine whether 6G6 would recognize Tyr39 nitration within longer αSyn fragments, and whether it would display selectivity toward the Tyr39 as compared to other αSyn tyrosine residues. These *in vitro* assays were performed using ELISA with electrochemiluminescence (ECL) detection. To test reactivity and specificity in the context of longer αSyn fragments, 6G6 was used as a capture antibody reacting with 50-mer HαSyn peptides that were nitrated at tyrosine positions 39, 125, 133 or 136. 6G6 bound Tyr39-nitrated peptides with an EC₅₀ of 14.8 ng/mL. A very weak signal was detected after incubations with relatively high concentrations of Tyr133-nitrated 50-mers, whereas no binding occurred between 6G6 and either Tyr125- or Tyr136-nitrated peptides (Fig. 1A). No binding was also detected after incubations in which non-nitrated Tyr39-containing 50-mers or unmodified full-length HαSyn were added to 6G6-coated plate wells (Fig. 1A).

**Figure 1.**
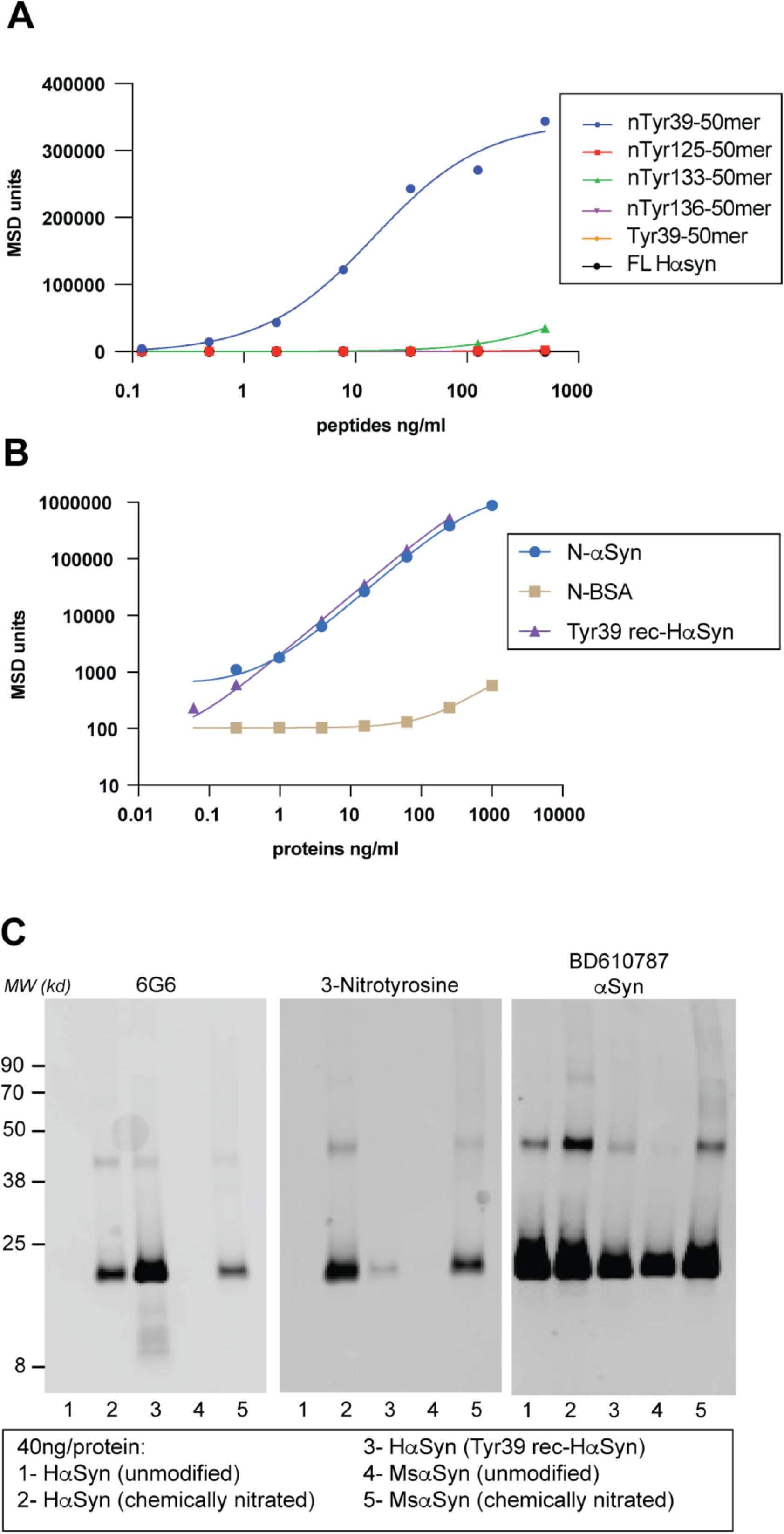
6G6 selectively recognizes Tyr39-nitrated αSyn. **(A)** Binding of 6G6 to biotinylated HαSyn 50-mer peptides nitrated at Tyr39 (nTyr39-50mer), Tyr125 (nTyr125-50mer), Tyr133 (nTyr133-50mer) or Tyr136 (nTyr136-50mer), as measured by electrochemiluminescent ELISA. Non-nitrated Tyr39 peptide (Tyr39-50mer) and unmodified full-length HαSyn (FL-HαSyn) were included as controls. **(B)** Binding of 6G6 to chemically nitrated full-length human α-synuclein (N-HαSyn), recombinant site-specifically Tyr39-nitrated full-length HαSyn (nTyr39 rec-HαSyn), or peroxynitrite-modified bovine serum albumin (N-BSA). **(C)** Western blot analysis showing the reactivity of 6G6, anti-3-nitrotyrosine, and total α-synuclein antibody (BD610787 αSyn) toward unmodified HαSyn, chemically nitrated HαSyn, recombinant site-specifically Tyr39-nitrated HαSyn, unmodified MsαSyn, and chemically nitrated MsαSyn.

To test antibody reactivity with full-length αSyn and rule out recognition of nitrated epitopes on unrelated proteins, specific reagents were generated. First, HαSyn was chemically nitrated using peroxynitrite; this treatment results in partial nitration at Tyr39 as well as nitration at other αSyn tyrosine sites.^23,30^ Furthermore, full length recombinant HαSyn (rec-HαSyn) nitrated at Tyr39 was produced by site-specific incorporation of nitrotyrosine as a genetically encoded “non-canonical” amino acid.^31^ Finally, nitrated bovine serum albumin (BSA) was obtained after treatment of the unmodified protein with peroxynitrite. In a direct binding ELISA, plates were coated with chemically nitrated HαSyn, site specifically nitrated rec-HαSyn or peroxynitrite-modified BSA. The coated plates were then incubated with 6G6, followed by detection with a species-specific SULFO-TAG antibody. 6G6 bound robustly both chemically nitrated HαSyn and Tyr39-nitrated rec-HαSyn, but showed no or very weak binding to chemically nitrated BSA even at very high concentrations (Fig. 1B). The binding properties of 6G6 were then further assessed by Western blot analyses that tested its reactivity against unmodified HαSyn, chemically nitrated HαSyn, site specifically nitrated rec-HαSyn, unmodified mouse αSyn (MsαSyn) and chemically nitrated MsαSyn. Data showed that 6G6 exclusively detected the nitrated forms of proteins, namely chemically nitrated human and mouse αSyn and Tyr39-nitrated rec-HαSyn (Fig. 1C, left panel). In control experiments, Western blots of unmodified and modified αSyn were analyzed using antibodies against either 3-nitrotyrosine (3-NT) or αSyn. As expected, anti-3-NT only reacted against nitrated αSyn forms whereas the anti-αSyn antibody detected all 5 unmodified and modified αSyn proteins (Fig. 1C, middle and right panels). It is noteworthy that, while both anti-3-NT and 6G6 detected Tyr39-nitrated rec-HαSyn, 6G6 reactivity was markedly more pronounced, underscoring its high sensitivity and specificity (Fig. 1C).

The *in vivo* experiments planned for this study involved administration of nitrated αSyn antibodies to mice. For this reason, the original rabbit 6G6 antibody was engineered and produced as a mouse chimeric IgG. To confirm that the chimeric antibody preserved properties of the parental rabbit clone, binding kinetics, affinities and specificity of rabbit and mouse 6G6 were compared using bio-layer interferometry (BLI; Octet platform) (Table 1). The binding affinity in the standard format BLI was determined to be 4.65nM and 4.29nM for the rabbit and mouse 6G6 antibodies, respectively (Table 1). Specific binding of 6G6 to Tyr39-nitrated αSyn was confirmed with inverted direct binding that showed no detectable reaction with Tyr125-, Tyr133- or Tyr136-nitrated αSyn peptides (data not shown). In the inverted BLI format mimicking multivalent binding and avidity effects, affinity values were below the lower detection limit of the assay (<10 pM). Taken together, these results demonstrate nanomolar to sub-nanomolar affinities of both rabbit and mouse-chimeric 6G6 to nitrated αSyn. They also reveal a positional specificity that, with either full-length αSyn or αSyn peptides, involves binding of 6G6 to HαSyn and MsαSyn at the same epitope site containing nitrated Tyr39.

**Table 1.**
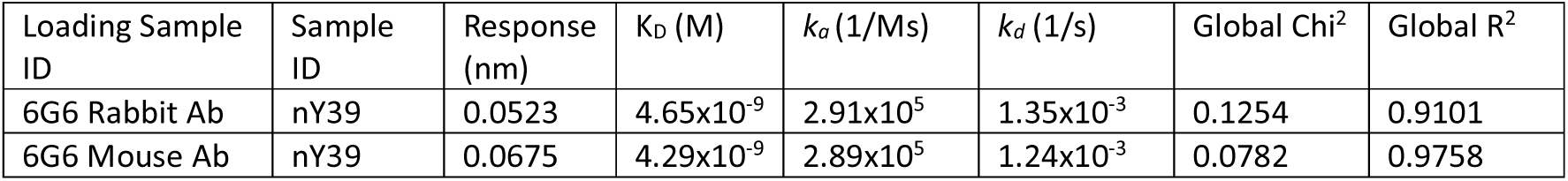
Kinetic characterization of rabbit and mouse-chimeric 6G6 binding to the Tyr39-nitrated αSyn 50-mer peptide.

### The anti-Tyr39-nitrated αSyn antibody, 6G6, counteracts HαSyn spreading in the paraquat/AAV vagal model

To test the hypothesis that Tyr39-nitrated αSyn plays a key role in αSyn interneuronal spreading, the effects of the anti-Tyr39-nitrated αSyn antibody, 6G6, were evaluated using the mouse AAV vagal model. In this model, when a vagal HαSyn-AAV injection triggers protein overexpression in the DMnX and NTS, caudo-rostral spreading of HαSyn is significantly enhanced by systemic administration of the herbicide paraquat (Musgrove et al., 2019). For the present study, mice were first all injected with HαSyn-AAVs into the vagus nerve and then treated with: (i) saline (instead of paraquat) and PBS (instead of antibodies), (ii) paraquat and PBS, (iii) saline plus IgG control, (iv) paraquat plus IgG control, (v) saline plus anti-Tyr39-nitrated αSyn (6G6), or (vi) paraquat plus 6G6 (Fig. 2A). Two single intraperitoneal injections of paraquat or saline were administered at 2 and 3 weeks after HαSyn-AAV treatment; mice in the IgG or 6G6 groups also received 4 weekly intraperitoneal injections of the antibody starting at 5 days post-AAVs. The entire treatment protocol was executed over a 4-week period, and animals were sacrificed at a time point that coincided with 1 week after the second paraquat exposure and 2 days after the last antibody administration (Fig. 2A). Levels and consistency of the 6G6 treatment were verified by MSD measurements of its plasma concentration at sacrifice. The antibody was detected in all treated animals, with similar terminal levels in the paraquat- and saline-injected groups (3050 ± 569 and 2677 ± 722 µg/mL, respectively; mean ± SD).

**Figure 2.**
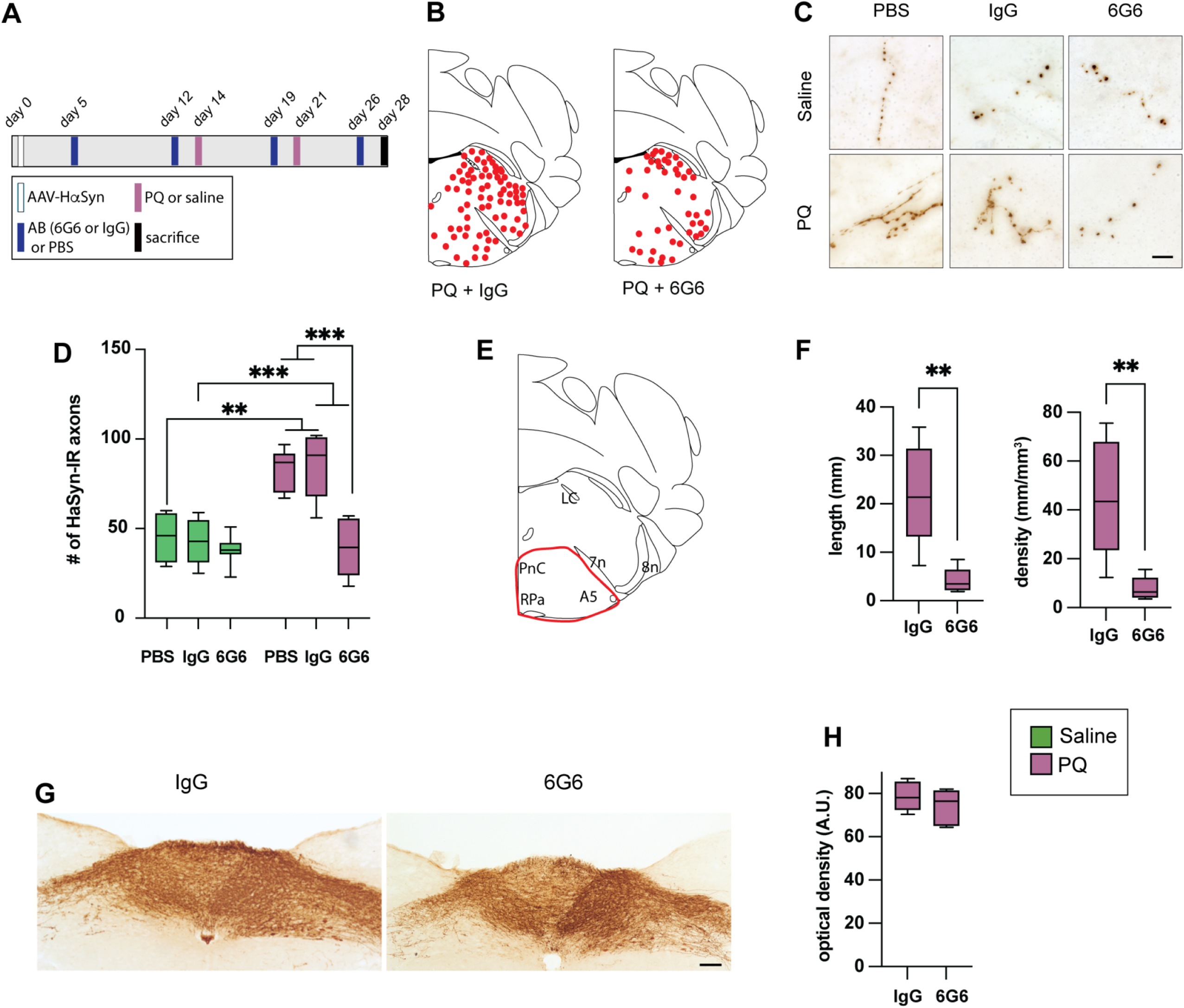
Anti-Tyr39-nitrated αSyn reduces HαSyn spreading in the paraquat/AAV vagal model. **(A)** Experimental design. Mice received a single unilateral injection of HαSyn-carrying AAVs into the left vagus nerve on day 0. Antibody treatment with control IgG or 6G6 was started 5 days later and continued once weekly for 4 weeks. Paraquat (PQ) or saline was administered intraperitoneally at 2 and 3 weeks after AAV injection. Animals were sacrificed 4 weeks after AAV injection. Coronal brain sections were stained with a specific HαSyn antibody. **(B)** Plots of HαSyn spreading in the pons obtained from PQ + IgG- and PQ + 6G6-treated mice. Red dots indicate the location of exogenous HαSyn-immunoreactive axons in the pons. **(C)** Representative pontine sections immunostained with anti-HαSyn, showing HαSyn-immunoreactive axons in the left pons. Scale bar = 5 µm. **(D)** HαSyn-immunoreactive axons were counted in the left (injected side) hemisphere of coronal pontine sections (n=4-8/group). Box and whisker plots show median, upper and lower quartiles, and maximum and minimum as whiskers. \*\**P* < 0.01; \*\*\**P* < 0.001, two-way ANOVA followed by Tukey post-hoc test. **(E-F)** Unbiased analyses of HαSyn-immunoreactive axons were made in a pontine area encompassing the ventral pons. **(E)** Illustration of the ventral pontine region used for stereological analysis. **(F)** Axonal length and density of HαSyn-immunoreactive axons estimated using the SpaceBalls stereological probe in PQ + IgG- and PQ + 6G6-treated mice (n=5/group). Box and whisker plots show median, upper and lower quartiles, and maximum and minimum as whiskers. \*\**P* < 0.01; unpaired *t*-test. **(G-H)** Assessment of HαSyn expression in the dorsal MO of AAV-injected mice. **(G)** Representative coronal sections of the medulla oblongata from PQ-treated mice receiving IgG or 6G6, immunostained with anti-HαSyn. Scale bar = 100 µm. **(H)** Densitometric analysis of medullary HαSyn staining in PQ + IgG- and PQ + 6G6-treated mice (n=5/group). Box-and-whisker plots show the median (middle line), upper and lower quartiles, and maximum and minimum values as whiskers.

Interneuronal spreading in the mouse AAV vagal model first and predominantly affects pontine regions. Therefore, to compare spreading in the presence or absence of paraquat as well as in IgG- *vs.* 6G6-treated mice, coronal sections of the pons were stained with anti-HαSyn and used for morphological and quantitative assessment of immunoreactive axons. Sections from all the AAV-injected animals were characterized by the presence of HαSyn-immunoreactive axons that appeared as winding threads with densely labeled and irregularly spaced varicosities. The distribution of these axons followed a pattern previously described for this vagal model (Helwig et al., 2016). In particular, HαSyn-containing fibers occupied pontine regions directly connected to the dMO, including the locus coeruleus and subcoeruleus nucleus, the parabrachial nuclei and pontine reticular areas (Fig. 2B). The density of HαSyn-stained fibers was overtly increased in samples from mice treated with paraquat/PBS or paraquat/IgG as compared to saline/PBS or saline/IgG. Quite in contrast, no significant changes were induced by paraquat exposure when the herbicide was administered in combination with 6G6 (Fig. 2C). Quantification of HαSyn-immunoreactive axons in the left (ipsilateral to the HαSyn-AAV injection) pons confirmed these microscopy observations. Axonal counts were significantly enhanced in mice treated with paraquat, regardless of whether they also received PBS or control IgG (Fig. 2D). This effect was abolished by treatment with anti-Tyr39-nitrated αSyn, since the number of pontine HαSyn-labeled axons was found to be comparable between saline- and paraquat-injected mice in the presence of 6G6 (Fig. 2D). Support for a robust protection of 6G6 against protein spreading was provided by analyses of pontine sections using the Spaceballs stereological probe. This tool allowed unbiased measurements of the length and density of HαSyn-labeled fibers in a predetermined, fixed area of the ventral pons (Fig. 2E). Comparisons were made between samples from mice treated with paraquat plus either IgG or 6G6, and data revealed a marked decrease in length and density measurements after administration of the anti-Tyr39-nitrated αSyn antibody (Fig. 2F).

Protein spreading in the AAV vagal model is significantly affected by the degree of HαSyn overexpression in the dMO (Helwig et al., 2022). Specific analyses were therefore carried out to verify that pattern and levels of HαSyn expression were comparable among the different treatment groups used for this study. When coronal sections of the MO were stained with anti- HαSyn, microscopy examination revealed robust immunoreactivity in the left (ipsilateral to the AAV injection) DMnX as well as the left and right NTS of all animals (Fig. 2G). This pattern of overexpression is consistent with a targeted transduction of DMnX neurons and neurons in the vagal ganglia after a unilateral vagal injection of HαSyn-AAVs (Kalia and Sullivan, 1982; Leslie et al., 1982; Pinto-Costa et al., 2023). The intensity of HαSyn immunoreactivity appeared to be similar across all treatment groups. In particular, both microscopy observation and semiquantitative densitometric analysis of stained medullary sections from paraquat/IgG- and paraquat/6G6-treated mice showed similar levels of HαSyn expression (Fig. 2G and H).

### 6G6 administration prevents spreading in transgenic mice with a *Gba1* mutation

Mutations of GBA1, the gene encoding the lysosomal enzyme glucocerebrosidase (GCase), are common genetic risk factors for PD (Sidransky et al., 2009; Vieira and Schapira, 2022). Enhanced neuronal vulnerability to oxidative/nitrative stress and increased HαSyn spreading have been reported in transgenic mice carrying a heterozygous L444P mutation of *Gba1* (L444P/wt mice) after a vagal injection with HαSyn-AVVs (La Vitola et al., 2024). A second set of experiments was therefore carried out to interrogate whether 6G6 administration may also affect HαSyn spreading in this GBA/vagal model. Spreading was compared in control wild-type (wt/wt) *vs.* L444P/wt mice injected with PBS and then assessed in transgenic animals treated with either IgG or 6G6. All mice received a vagal AAV injection of HαSyn-AVVs. The subsequent treatment followed a schedule summarized in Fig. 3A that included 4 weekly intraperitoneal injections with PBS or antibodies. In the group of animals treated with 6G6, dosage was verified by detection of the antibody in plasma specimens collected at the time of sacrifice; terminal 6G6 concentrations were found to be 3368 ± 468 µg/mL (mean ± SD).

**Figure 3.**
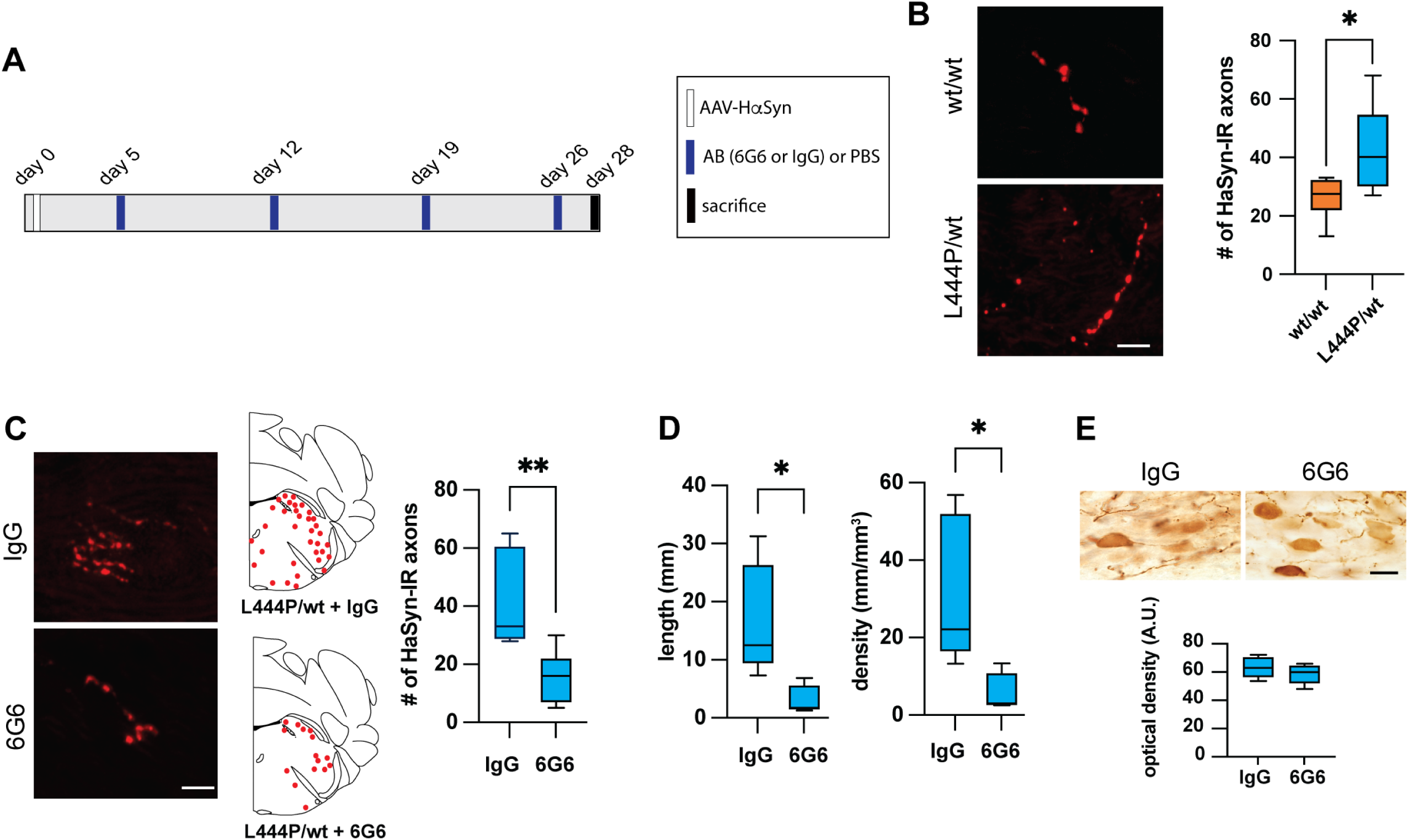
Anti-Tyr39-nitrated αSyn reduces HαSyn spreading in Gba1L444P/wt mice. **(A)** Experimental design. Wild-type (wt/wt) and heterozygous Gba1L444P/wt mice received a single unilateral injection of HαSyn-carrying AAVs into the left vagus nerve. Antibody treatment with control IgG or 6G6 was started 5 days later and continued once weekly for 4 weeks. Animals were sacrificed 4 weeks after AAV injection. Coronal brain sections were stained with a specific HαSyn antibody. **(B-D)** Pons sections were immunostained with anti-HαSyn. **(B)** AAV-mediated HαSyn expression and quantification of HαSyn-immunoreactive axons in the left (injected side) pons comparing wt/wt and L444P/wt mice treated with PBS (n = 6/group). Box-and-whisker plots show the median, upper and lower quartiles, and maximum and minimum values as whiskers. *P* < 0.05, unpaired *t*-test. Scale bar = 5 µm. **(C)** HαSyn-immunoreactive axons in pontine sections of L444P/wt mice treated with IgG or 6G6 and distribution of exogenous HαSyn-immunoreactive axons in the pons. HαSyn-immunoreactive axons were counted in the left (injected side) hemisphere of coronal pontine sections in IgG- and 6G6-treated L444P/wt mice (n = 6 and 7, respectively). Box-and-whisker plots show the median, upper and lower quartiles, and maximum and minimum values as whiskers. **\*\****P* < 0.01, unpaired *t*-test. Scale bar = 5 µm. **(D)** Stereological measurements of HαSyn-positive axonal length and density in the pons of IgG- and 6G6-treated L444P/wt mice (n=5/group) using the SpaceBalls probe. Box-and-whisker plots show the median, upper and lower quartiles, and maximum and minimum values as whiskers. \**P* < 0.05, unpaired *t*-test. **(E)** Representative medullary sections immunostained with anti-HαSyn, showing comparable levels of AAV-mediated transgene expression in the DMnX of IgG- and 6G6-treated L444P/wt mice (n=5/group), together with densitometric quantification of HαSyn staining in medullary sections from the two groups. Scale bar = 20 µm. Box-and-whisker plots show the median, upper and lower quartiles, and maximum and minimum values as whiskers.

At the time of sacrifice, brains were removed and fixed in paraformaldehyde. Coronal sections of the pons were immunostained with anti-HαSyn for quantification of HαSyn-containing axons as markers of medullary-to-pontine protein spreading. In line with earlier results (La Vitola et al., 2024), manual counting showed a significant increase in the number of labeled axons in L444P/wt as compared to wt/wt animals injected with PBS (42.67 ± 15.11 vs 26.33 ± 7.20, mean ± SD; Fig. 3B). Analyses then focused on HαSyn spreading in transgenic mice treated with either IgG or 6G6 and, for these analyses, counting of pontine immunoreactive axons was paralleled by measurements of their length and density. Microscopy observation already revealed that the density of labeled axons was markedly reduced in the group of mice receiving 6G6, and manual counting determined that the number of HαSyn-containing axons was substantially decreased by **>**2.5 fold (from 41.2 ± 16.4 to 15.1 ± 8.8, mean ± SD) after treatment with the anti-Tyr39-nitrated αSyn antibody (Fig. 3C). Similarly, measurements using the Spaceballs tool found values of axonal length and density that were reduced by 81.1% and 81.2%, respectively, decreasing from 16.79 ± 9.57 to 3.17 ± 2.40 and from 31.78 ± 18.99 to 5.97 ± 4.76 (mean ± SD; Fig. 3D). The effect of 6G6 on medullary-to-pontine protein spreading could not simply be attributed to differences in AAV-induced HαSyn overexpression between IgG- *vs.* 6G6-treated L444P/wt mice. This possibility was in fact ruled out by microscopy observations and densitometric measurements on coronal MO sections stained with anti- HαSyn; these analyses confirmed comparable levels of HαSyn transduction regardless of the antibody treatment (Fig. 3E).

Taken together with the results using the paraquat/AAV vagal model, these data support a direct involvement of nitrated αSyn in pathological spreading processes that can indeed be effectively mitigated by antibody blockade intervention with anti-Tyr39-nitrated αSyn.

### 6G6 protects against PFF-induced P-αSyn pathology and nigral neuronal loss

To further assess the efficacy of 6G6 in an α-synucleinopathy model that does not rely on viral overexpression, a final set of experiments was carried out in homozygous M83 transgenic mice expressing human A53T mutant αSyn under the Prnp promoter. In these mice, an intrastriatal injection of αSyn PFFs triggers robust P-αSyn pathology that spreads throughout the brain; in the substantia nigra, accumulation of this pathology is accompanied by frank degeneration of dopaminergic neurons (Luk et al., 2012). Young adult M83 mice were given a single unilateral injection of 4 µg HαSyn PFFs into the dorsal striatum; control animals were injected with an equivalent amount of monomeric αSyn. Antibody treatment (IgG or 6G6) was initiated 2 hours prior to PFF inoculation and continued once a week for the following 7 weeks (Fig. 4A). Terminal 6G6 plasma concentrations were 2042 ± 336 µg/mL (mean ± SD). 6G6 levels were also measured in olfactory bulb tissue dissected post-mortem and found to be 5.65 ± 1.34 µg/mL (mean ± SD). The resulting brain-to-plasma ratio was 0.28%, consistent with earlier data on CNS exposure after systemic administration of a monoclonal antibody.^32^

**Figure 4.**
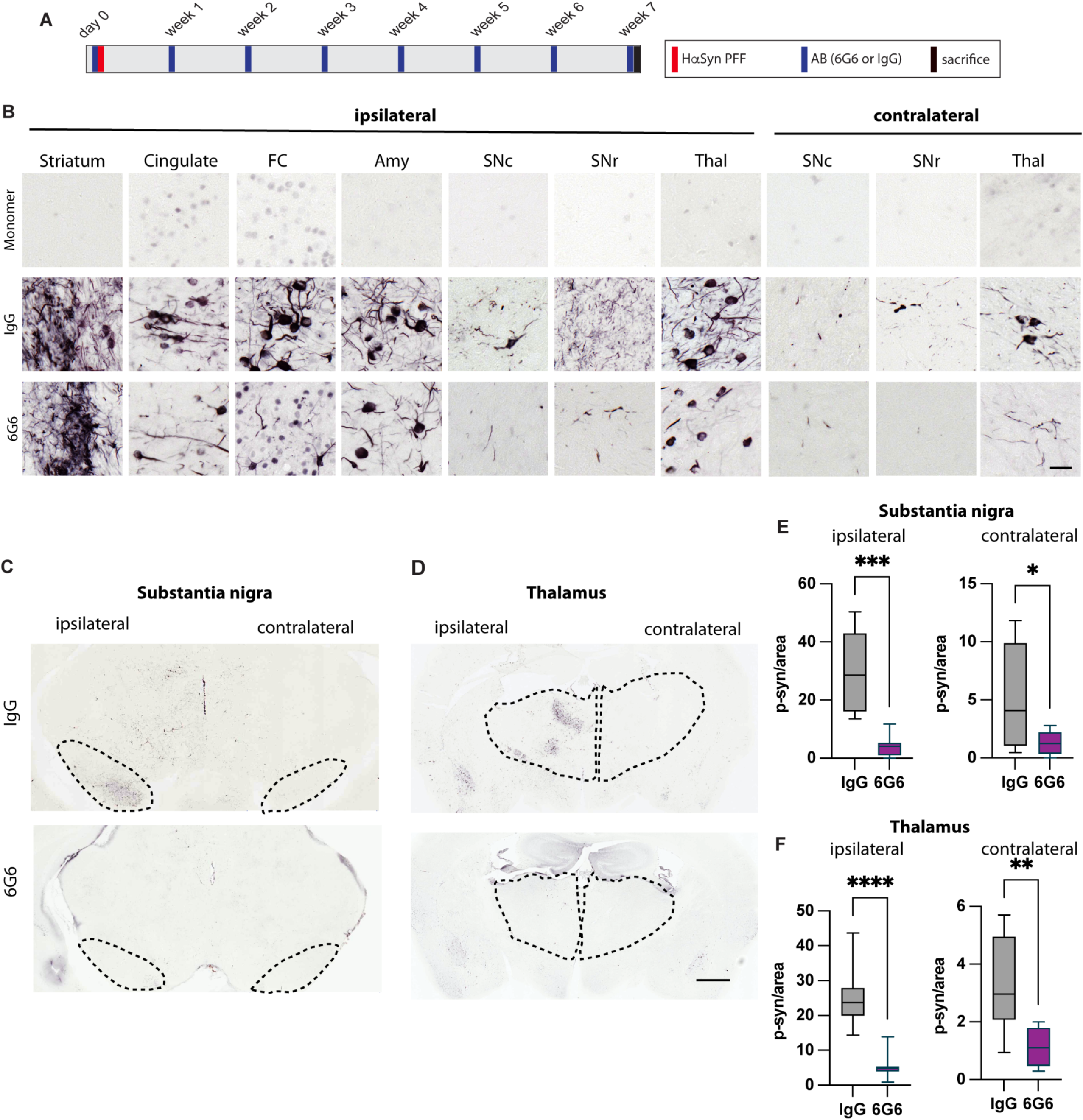
Anti-Tyr39-nitrated αSyn reduces HαSyn spreading in M83 PFF model. **(A)** Experimental design. Homozygous M83 mice received a single unilateral intrastriatal injection of HαSyn PFFs or monomer on day 0. Antibody treatment with control IgG or 6G6 was started 2 h before inoculation and continued once weekly for 7 weeks. Animals were sacrificed at study week 7, 2 days after the final antibody injection. Coronal brain sections were immunostained with a specific antibody against P-αSyn. **(B)** Representative images from coronal sections immunostained for P-αSyn, showing pathology in the striatum, cingulate cortex, frontal cortex (FC), basolateral amygdala (Amy), substantia nigra pars compacta (SNc), substantia nigra pars reticulata (SNr), and thalamus ipsilateral to the injection side, as well as in the SNc, SNr and thalamus contralateral to the injection side. Scale bar = 50 µm. **(C, D)** Low-magnification images showing the regions of interest used for P-αSyn quantification in the substantia nigra and thalamus. Scale bar = 1 mm. **(E)** Quantification of P-αSyn-positive objects per analysed area in the substantia nigra ipsilateral and contralateral to the injection side in IgG-treated (n = 9) and 6G6-treated (n = 8) mice. Box-and-whisker plots show the median, upper and lower quartiles, and maximum and minimum values as whiskers. *P* < 0.05; \*\**P* < 0.001, unpaired *t*-test. **(F)** Quantification of P-αSyn-positive objects per analysed area in the thalamus ipsilateral and contralateral to the injection side in IgG-treated (n = 9) and 6G6-treated (n = 8) mice. Box-and-whisker plots show the median, upper and lower quartiles, and maximum and minimum values as whiskers. \*\**P* < 0.001, \*\*\*\**P* < 0.0001; unpaired *t*-test.

Brains were collected post-mortem from all PFF-treated M83 mice. They were then cut into coronal sections that were stained with anti-P-αSyn. Microscopy examination showed widespread P-αSyn pathology throughout basal ganglia and extra–basal ganglia regions that included the dorsal striatum, substantia nigra pars compacta (SNc), substantia nigra pars reticulata (SNr), thalamus, amygdala and cortex (Fig. 4B). P-αSyn-immunoreactive lesions were evident bilaterally but markedly more pronounced on the side of the brain ipsilateral to the PFF injection; images in Fig. 4B-D illustrate this lateralization in sections from the SN and thalamus, two regions severely affected by P-αSyn accumulation. Microscopy analysis also compared the pathology burden in mice treated with IgG *vs.* 6G6 and revealed an overt reduction of both neurites and cell bodies immunoreactive for P-αSyn after administration of the anti-Tyr39-nitrated αSyn antibody (Fig. 4B-D). Quantification of P-αSyn-positive neurites and cell bodies in sections from the SN and thalamus confirmed this significant effect of 6G6. In the ipsilateral SN, mice receiving IgG showed 28.8 ± 14.1 objects/area (mean ± SD), whereas mice injected with 6G6 had 4.2 ± 3.6 objects/area (85% reduction) (Fig. 4C,E). In the contralateral SN, measurements showed a 75% pathology reduction in 6G6- *vs.* IgG-treated mice, with values being 1.3 ± 1.0 and 5.1 ± 4.4 objects/area, respectively (Fig. 4C,E). Quantitative assessment of thalamic sections yielded similar results. Sections from the ipsilateral and contralateral thalamus of IgG-injected mice displayed 24.8 ± 8.3 and 3.3 ± 1.6 objects/area, as compared to samples from 6G6-treated animals containing 5.4 ± 3.7 (78% reduction) and 1.1 ± 0.6 (67% reduction) objects/area (Fig. 4D,F).

The next set of analyses assessed whether, as previously reported (Luk et al., 2012), injections of HαSyn PFFs into the striatum of M83 mice resulted in nigral dopaminergic cell death and whether, using this experimental paradigm, administration of 6G6 affected the integrity/survival of nigral neurons. Coronal midbrain sections were both immunolabeled with anti-tyrosine hydroxylase (TH, a marker of dopaminergic cells) and Nissl-stained and used for microscopy examination and unbiased stereological neuronal counting. When compared to microscopy observations in control mice, an overt reduction of nigral TH immunoreactivity and nigral TH-positive neurons characterized the SN of PFF-injected animals treated with IgG (Fig. 5A). This effect contrasted to observations in midbrain sections from mice treated with PFFs and 6G6 in which TH immunoreactivity was preserved and the density of TH-labeled cells appeared comparable to neuronal density in the SN of control animals (Fig. 5A). Unbiased stereological counting of TH-positive neurons was carried out in the right (ipsilateral to the PFF injection side) SNc to verify and precisely quantify treatment-associated differences in dopaminergic cell number. Similar to data reported in naïve wild-type mice ^33^, nigral cell counts in monomer-injected M83 mice were 6910 ± 75 (mean ± SD) (Fig. 5B). PFF treatment in combination with IgG injections caused a significant loss of TH-positive cell, with count values declining to 4743 ± 864 (-31%) (Fig. 5B). Using the same PFF treatment but together with 6G6 administration, the number of nigral dopaminergic neurons was found to be 6132 ± 913. This value was not statistically different than the cell number in control mice; it was instead significantly higher than the count of TH-positive neurons in PFF/IgG-treated animals (Fig. 5B). Treatment-induced changes in the number of TH-labeled cells may not necessarily be interpreted as evidence of neuronal loss/survival, since they could also arise from phenotypic differences in TH regulation. For this reason, nigral neurons in control and PFF-injected mice were also stereologically counted using their Nissl staining for cell identification. Counts yielded 9333 ± 236 (mean ± SD), 6624 ± 976 (-29% as compared to the control value) and 7975 ± 810 (not statistically different from the control value) Nissl-positive neurons in monomer-injected, PFF/IgG-treated and PFF/6G6-administered animals, respectively (Fig. 5C). Thus, taken together, counts of both TH-immunoreactive and Nissl-stained cells clearly indicate that PFF-induced nigral αSyn pathology was accompanied by degeneration of dopaminergic cells and that this neuronal loss could be largely prevented by anti-Tyr39-nitrated αSyn administration.

**Figure 5.**
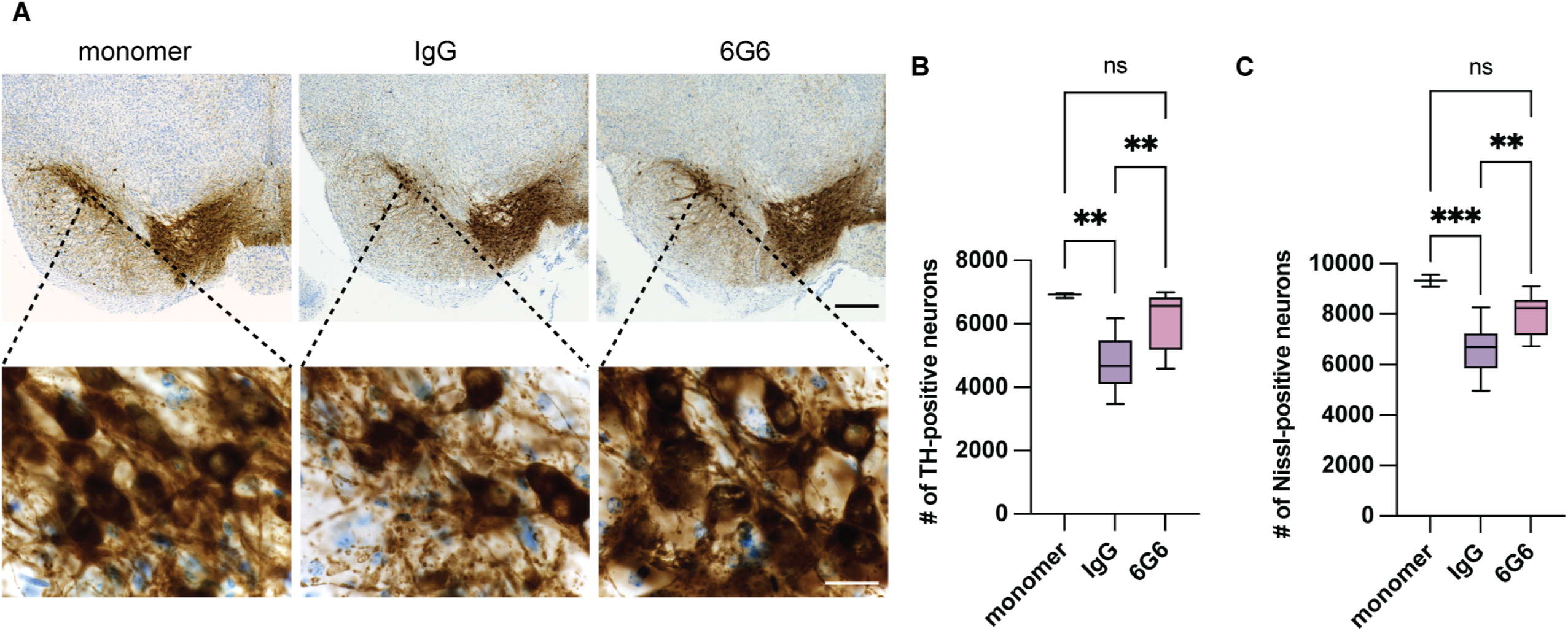
Anti-Tyr39-nitrated αSyn preserves nigral neurons in PFF-injected M83 mice. Coronal midbrain sections from monomer-injected (n = 3) control mice and PFF-injected mice treated with either IgG (n = 10) or 6G6 (n = 9) were stained for tyrosine hydroxylase (TH, brown stain) and counterstained with Nissl (blue stain). **(A)** Low-magnification images (upper panels) and high-magnification images (lower panels) of representative midbrain sections containing the substantia nigra pars compacta (SNc). Scale bars: upper panels, 100 µm; lower panels, 20 µm. **(B)** Unbiased stereological quantification of TH-positive neurons in the ipsilateral SNc of monomer-injected, IgG-treated, and 6G6-treated mice. Box-and-whisker plots show the median, upper and lower quartiles, and maximum and minimum values as whiskers. *P <* 0.01; *\*\*P* < 0.001, one-way ANOVA followed by Tukey’s post hoc test. **(C)** Unbiased stereological quantification of Nissl-positive neurons in the ipsilateral SNc of monomer-injected, IgG-treated, and 6G6-treated mice. Box-and-whisker plots show the median, upper and lower quartiles, and maximum and minimum values as whiskers. *\*\*P* < 0.01; *\*\*\*P* < 0.001, one-way ANOVA followed by Tukey’s post hoc test.

### Tyr39-nitrated αSyn is elevated in CSF samples from PD patients

The findings described above point to an important role of nitrated αSyn in experimental models of PD-like pathological processes. To assess and support the translational relevance of these findings, a pilot study was designed to interrogate whether levels of Tyr39-nitrated αSyn could be detected in human CSF samples and, if so, whether they would be elevated in specimens from PD patients. A highly sensitive assay was developed using the Quanterix SIMOA platform; for this assay, the 8C2 anti-Tyr39-nitrated αSyn antibody served as capture reagent whereas a total anti-αSyn antibody was used for detection purposes. The limit of Tyr39-nitrated αSyn quantification was 0.01 pg/mL.

CSF samples, which were collected from 10 PD patients and 5 sex- and age-matched individuals with no signs/symptoms of PD (Table 2), were assayed for Tyr39-nitrated αSyn as well as total protein and total αSyn concentrations. Total protein levels were 528.2 ± 145 (mean ± SD) and 658.8 ± 263.9 µg/mL, and total αSyn concentrations were 482.7 ± 123.2 and 438.7 ± 137.0 pg/mL in non-PD and PD specimens, respectively, indicating lack of significant disease-associated differences (Fig. 6A,B). Quite in contrast, measurements of Tyr39-nitrated αSyn yielded values that clearly separated patients from non-patient individuals and are consistent with enhanced nitrative burden associated with the disease status. Tyr39-nitrated αSyn was very low (0.07 ± 0.07 pg/mL) in non-PD CSF and, in fact, did not reach the detection limit in samples from 3 of the non-patient individuals. It instead was present in the CSF of all PD patients, with values ranging from 3.2 to 7.1 pg/mL and an average concentration of 4.9 ± 1.3 pg/mL (Fig. 6C).

**Figure 6.**
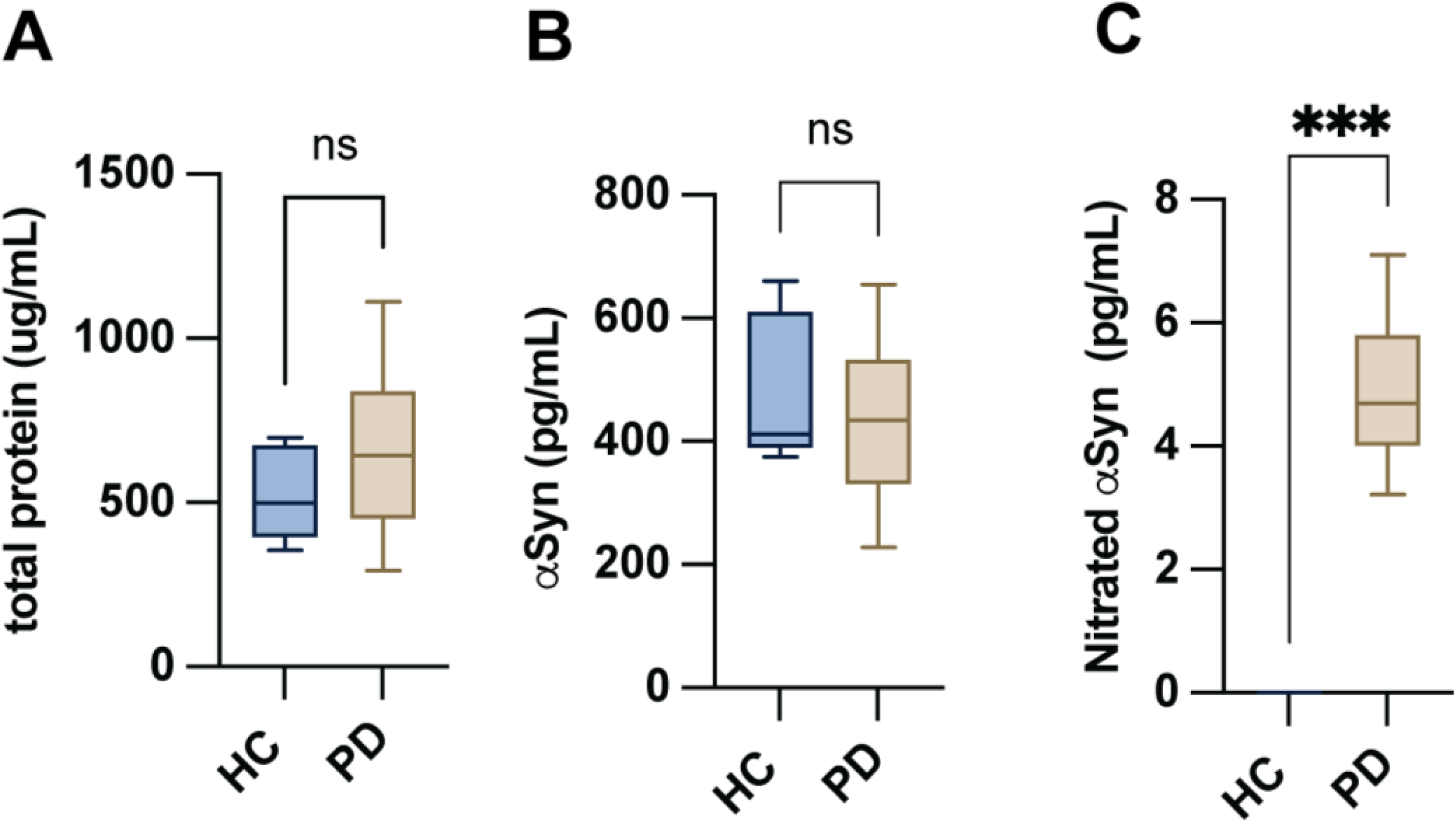
Tyr39-nitrated αSyn in human cerebrospinal fluid. **(A)** Total protein concentrations in cerebrospinal fluid samples from healthy (n=5) controls and Parkinson’s disease (n=10) patients. **(B)** Total αSyn concentrations in the same CSF samples. **(C)** Tyr39-nitrated αSyn concentrations measured by SIMOA in CSF samples from healthy controls and Parkinson’s disease patients. Box-and-whisker plots show the median, upper and lower quartiles, and maximum and minimum values as whiskers. \*\*\**P* < 0.001, unpaired *t*-test.

**Table 2.**
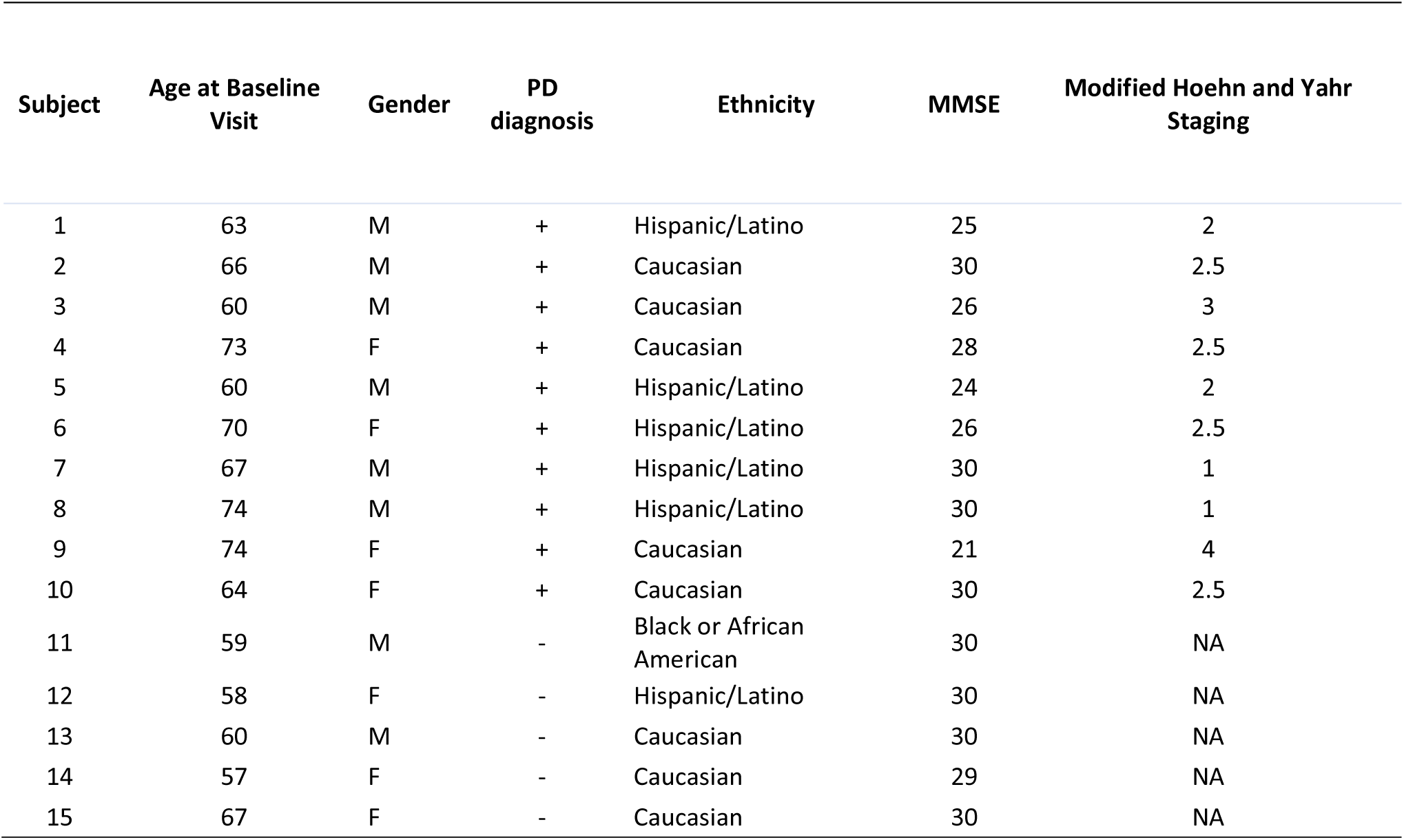
Demographic and clinical characteristics of the human CSF sample cohort.

| **Subject** | **Age at Baseline Visit** | **Gender** | **PD diagnosis** | **Ethnicity** | **MMSE** | **Modified Hoehn and Yahr Staging** |
| --- | --- | --- | --- | --- | --- | --- |
| 1 | 63 | M | + | Hispanic/Latino | 25 | 2 |
| 2 | 66 | M | + | Caucasian | 30 | 2.5 |
| 3 | 60 | M | + | Caucasian | 26 | 3 |
| 4 | 73 | F | + | Caucasian | 28 | 2.5 |
| 5 | 60 | M | + | Hispanic/Latino | 24 | 2 |
| 6 | 70 | F | + | Hispanic/Latino | 26 | 2.5 |
| 7 | 67 | M | + | Hispanic/Latino | 30 | 1 |
| 8 | 74 | M | + | Hispanic/Latino | 30 | 1 |
| 9 | 74 | F | + | Caucasian | 21 | 4 |
| 10 | 64 | F | + | Caucasian | 30 | 2.5 |
| 11 | 59 | M | - | Black or African American | 30 | NA |
| 12 | 58 | F | - | Hispanic/Latino | 30 | NA |
| 13 | 60 | M | - | Caucasian | 30 | NA |
| 14 | 57 | F | - | Caucasian | 29 | NA |
| 15 | 67 | F | - | Caucasian | 30 | NA |

## Discussion

Nitration of αSyn in Lewy bodies and Lewy neurites, the neuronal inclusions pathognomonic of PD, provides direct evidence that oxidative and nitrative reactions occur in cells bearing PD pathology.^18^ This observation is not only consistent with the view that nitrated αSyn is a marker of specific pathogenetic mechanisms. It also suggests that, by altering properties and behavior of αSyn, nitrative reactions could generate post-translationally modified protein species with distinct and potentially more pronounced pathogenic properties. In the present study, the occurrence and relevance of αSyn nitration in PD was underscored by the results of measurements of nitrated αSyn, in particular Tyr39-nitrated αSyn, in human CSF samples that revealed its marked increase in patients as compared to non-PD controls. In its experimental component, this study also provided strong evidence that nitrated αSyn formation is directly linked to deleterious consequences in animal models characterised by CNS spreading of αSyn pathology. Finally, our current results indicate that targeting nitrated αSyn burden is feasible To investigate the pathogenic role of Tyr39-nitrated αSyn and assess the consequences of its blockade, a new antibody, named 6G6, was specifically designed and charactezized. In a series of *in vitro* assays, 6G6 showed high-affinity binding to nitrated αSyn with positional specificity for Tyr39 and negligible reactivity toward non-nitrated αSyn or αSyn nitrated at its other tyrosine residues. Both rabbit 6G6 and its chimeric mouse antibody bound the same αSyn epitope and displayed similar biochemical properties; the latter antibody was then utilized for treatment of mice with well-defined spreading αSyn pathology. An important component of this study was its design aimed at testing the effects of 6G6 in three different *in vivo* models. The antibody was first administered to mice in which interneuronal HαSyn transfer and consequent caudo-rostral protein spreading were triggered by AAV-induced HαSyn overexpression in the medullary dorsal vagal system.^13,14,16^ In order to exacerbate HαSyn transfer from medullary donor neurons to pontine recipient axons, two strategies were adopted: AAV-injected mice were exposed to the herbicide paraquat or AAV-induced HαSyn overexpression was triggered in *Gba1* mutant mice.^17,26^ Paraquat injections and treatment of Gba1 transgenics were also chosen to mimic the effects that environmental exposures and genetic predisposition factors could have on brain nitrative reactions and ensuing PD-like pathology. A separate set of experiments was carried out in a model of αSyn pathology that did not involve vector-induced protein overexpression but was triggered by an intraparenchymal brain injection of αSyn PFFs. This widely used experimental paradigm is thought to cause αSyn aggregation and spreading through a prion-like mechanism by which seeds of misfolded αSyn act as templates that “corrupt” normal endogenous αSyn and thus propagate pathogenic protein conformations.^7,34^ When HαSyn PFFs are injected into the mouse striatum, as done in this study, aggregate pathology spreads to regions that include the thalamus and SN; in the latter, αSyn lesions are accompanied by evidence of neuronal injury and, ultimately, neurodegeneration.^7,35^

In the paraquat/AAV and GBA/AAV models, experiments compared medullary-to-pontine HαSyn spreading in the absence and presence of 6G6 administration whereas, in the PFF model, accumulation of P-αSyn-labeled lesions was assessed in the thalamus and SN and neurodegeneration quantified in the SNc of M83 mice treated with 6G6 or control IgG. Blockade of Tyr39-nitrated αSyn with 6G6 resulted in significant neuroprotective effects under all three experimental paradigms. Number, density and length of pontine axons containing HαSyn were significantly reduced in AAV-injected animals receiving 6G6 and, similarly, 6G6 administration markedly alleviated the PFF-induced burden of P-αSyn pathology in both thalamic and nigral mouse tissue sections. A loss of TH-immunoreactive and Nissl-stained neurons characterized the SNc of mice that received PFF injections and were treated with IgG while, quite noticeably, animals receiving PFFs and 6G6 showed a near-complete preservation of TH-positive cells and robust rescue of total Nissl-positive neurons. Thus, in parallel to preventing P-αSyn lesions, 6G6 preserved the integrity of nigral dopaminergic neurons. Taken together, these findings provide strong evidence in favor of a role of Tyr39-nitrated αSyn in PD-like pathogenetic processes and indicate that blockade of this modified αSyn species with targeted antibody treatment can effectively lead to neuroprotection.

The ability of 6G6 to counteract αSyn pathology under all three experimental paradigms used in this study supports a common sequence of toxic events that, through the formation and accumulation of reactive oxygen (ROS) and nitrative (RNS) species, result in nitrative αSyn modifications.^36^ In the paraquat/AAV model, ROS and RNS are likely to be generated as a result of redox cycling reactions that are catalyzed by neuronal as well as glial enzymes (e.g., nitric oxide synthase and NADPH oxidase) and involve one-electron transfers between paraquat and molecular oxygen.^37^ The occurrence of αSyn nitrative reactions in AAV-injected mice carrying the L444P *Gba1* mutation is in line with the results of a recent investigation showing a relationship between neuronal expression of this mutation and mitochondrial oxidative/nitrative (Ox/Nt) stress.^17^ Finally, mitochondrial damage and Ox stress have been described as consequences of neuronal exposure to αSyn PFFs and suggested to act, at least in part, as cell-autonomous vulnerability factors for the propagation of PFF-induced P-αSyn lesions.^38,39^ Ox/Nt stress has long been implicated in PD pathogenesis.^36,40^ Results of this study showing a neuroprotective effect of 6G6 further support this mechanistic role. As importantly, however, they also highlight the generation of nitrated αSyn species as a critical link between Ox/Nt stress and the development and spreading of αSyn pathology.

Two other important considerations are raised by the effectiveness of 6G6 in counteracting αSyn spreading pathology. The first relates to the mechanism(s) of action of this antibody treatment. Earlier in vitro and in vivo work has demonstrated that αSyn nitration generates protein species with enhanced intercellular mobility and pro-aggregating properties.^17,26,27^ Particularly relevant were the results of earlier experiments revealing that cell-to-cell αSyn transfer was significantly exacerbated by oxidative/nitrative stress and could be blocked by addition of anti-N-αSyn antibodies to cell culture incubation media.^26^ It is conceivable, therefore, that in the AAV and PFF models assessed in this study, highly mobile Tyr39-nitrated αSyn may represent a targetable vehicle for pathological αSyn spreading and a mediator of toxic mechanisms, such as prion-like αSyn templating and aggregation. The second consideration raised by findings of this study concerns the fact that, while indicating an important deleterious role of oxidative/nitrative stress and Tyr39-nitrated αSyn, they do not necessarily rule out the possibility that other αSyn species and other mechanisms may contribute to protein mobility and ensuing pathology. This possibility would be consistent with data showing that 6G6 significantly reduced but did not completely abolish αSyn spreading and aggregation. 6G6 specifically targets Tyr39-nitrated αSyn. Therefore, should other nitrated αSyn forms, namely Tyr125-, Tyr133- and Tyr136-nitrated αSyn, or other modified αSyn species also possess high mobility and pro-aggregating properties, they could play a role in the formation and spreading of pathological lesions even in the presence of 6G6 treatment. Furthermore, 6G6-resistant αSyn spreading pathology may arise from molecular mechanisms other than oxidative/nitrative stress that include lysosomal alterations, changes in neuronal activity, formation of tunneling nanotubes and enhanced αSyn release through exosomes and unconventional exocytosis.^27,41–44^ Further investigations are warranted to assess whether the protective effects of 6G6 could be enhanced by complementary treatments targeting other modified αSyn species and counteracting other mechanisms of αSyn burden and propagation.

From the therapeutic standpoint, it is important to note that, over the past several years, treatments with a variety of αSyn antibodies have been proposed and tested both experimentally and in clinical trials. These antibodies, including cinpanemab, prasinezumab, MEDI1341, amlenetug (Lu AF82422) and exidavnemab, were primarily designed and selected based on their ability to preferentially bind aggregated αSyn forms.^45–51^ Cinpanemab and prasinezumab were initially used in pre-clinical investigations and, then, also tested for their effectiveness in Phase II clinical trials. They both showed protective effects in animal models of αSyn aggregate pathology.^52,53^ However, a clinical study assessing cinpanemab treatment in PD patients failed to meet its primary and secondary end points.^45^ The prasinezumab Phase II investigation is still ongoing. Its initial results in early-stage PD patients showed lack of effectiveness in meeting the primary end point of the trial but also suggested efficacy in slowing disease motor progression.^46,47^ Taken together, current data on antibody-based treatment targeting αSyn raise a number of yet unanswered questions. One of these questions concerns the apparent discrepancy between significant findings in PD animal models and relatively unsatisfactory results of clinical investigations. Reason(s) of such discrepancy remain unclear. It has been suggested, however that discordant experimental *vs.* clinical outcomes may be explained, at least in part, by a distinct αSyn cleavage or other significant differences in post-translational αSyn modifications in the human brain.^54^ Alternatively, the magnitude of effect in PD animal models with prasinezumab and cinpanemab were small suggesting a need for a more substantial effect, such as that observed with this nitrated synuclein Ab, may be necessary to show stronger disease modifying effects in PD patients.

The results of earlier clinical investigations also highlight the need for further research on therapeutic strategies that would counteract αSyn pathology by targeting other (besides aggregated) “toxic” αSyn forms and specific mechanisms of toxic αSyn conversion. Findings of the present study provide *in vivo* proof-of-concept that blocking a disease-associated post-translational modification of αSyn may be sufficient to counteract and alleviate PD pathology. In particular, the protective effects of 6G6 against αSyn aggregation and spreading and against αSyn-induced neuronal injury identifies Tyr39-nitrated αSyn as a targetable mediator of toxic/pathological processes. As a corollary to the deleterious effects of nitrated αSyn, Ox/Nt reactions should be considered important mechanisms for toxic αSyn conversion and, as such, be themselves targeted for the development of new therapeutics. For example, based on recent work showing a catalytic role of GLOD4 in αSyn nitration^28,29^, inhibition of this enzyme may work upstream to prevent the generation of pro-aggregating and highly mobile nitrated αSyn species, thus reducing αSyn toxic burden. As a final consideration, future testing of therapeutic strategies against nitrated αSyn and/or Ox/Nt reactions should consider the possibility that effectiveness of these therapies may be more pronounced in subpopulations of patients with higher susceptibility to Ox/Nt stress. Results of the current and earlier investigations support a relationship between loss of GCase activity and increased ROS/RNS production^17,55^, thus suggesting that patients carrying *GBA1* mutations may be among the better candidates for anti-nitrated αSyn intervention.

### Antibody generation and engineering

Rabbit monoclonal antibodies against Tyr39-nitrated αSyn were generated at Abclonal (Woburn, MA, USA). Rabbits were immunised with a nitrated 9-mer peptide corresponding to human αSyn aa35–43 (EGVL(nitro-Y)VGSK) on days 0, 7, 21, 42 and 59. Complete Freund’s adjuvant was used on day 0 and incomplete Freund’s adjuvant on days 7, 21 and 42; peptide alone was administered on day 59. On day 63, spleens were collected and antibody-producing B cells were cultured.

Culture supernatants were screened by ELISA for specific binding to the nitrated immunogen and lack of binding to the non-nitrated peptide. Positive clones were selected for IgG gene cloning and small-scale expression. Top candidates were subcloned into expression vectors, expressed and purified for further characterisation.

Clone 8C2 was selected as an assay reagent and clone 6G6 as the lead candidate for in vivo studies. Both antibodies were engineered as mouse chimeric IgG1 antibodies (Absolute Antibodies, Cleveland, UK). Large-scale production of mouse chimeric 6G6 and 8C2 was carried out at Absolute Antibodies and ProBio (GenScript ProBio USA, Piscataway, NJ, USA).

### Peptides, recombinant proteins, modifications and pre-formed fibrils

The αSyn peptides and full-length human αSyn (Abclonal) were used for ELISA, affinity, and specificity studies. The sequences were as follows: αSyn 50-mer aa14–64, nY39: Biotin-GVVAAAEKTKQGVAEAAGKTKEGVL(nitrated-Y)VGSKTKEGVVHGVATVAEKTKEQVT; αSyn 50-mer aa14–64, Y39: Biotin-GVVAAAEKTKQGVAEAAGKTKEGVLYVGSKTKEGVVHGVATVAEKTKEQVT; αSyn 50-mer aa90–140, nY125: Biotin-AATGFVKKDQLGKNEEGAPQEGILEDMPVDPDNEA(nitrated-Y)EMPSEEGYQDYEPEA; αSyn 50-mer aa90–140, nY133: Biotin-AATGFVKKDQLGKNEEGAPQEGILEDMPVDPDNEAYEMPSEEG(nitrated-Y)QDYEPEA; αSyn 50-mer aa90–140, nY136: Biotin-AATGFVKKDQLGKNEEGAPQEGILEDMPVDPDNEAYEMPSEEGYQD(nitrated-Y)EPEA; αSyn 70-mer aa31–100, nY39: GKTKEGVL(nitrated-Y)VGSKTKEGVVHGVATVAEKTKEQVTNVGGAVVTGVTAVAQKTVEGAGSIAAAT GFVKKDQL. A non-nitrated 50-mer Y39 peptide (aa14–64; Biotin-GVVAAAEKTKQGVAEAAGKTKEGVLYVGSKTKEGVVHGVATVAEKTKEQVT) and biotinylated full-length human αSyn were also included.

For chemical nitration, a-synuclein protein (mouse and human, rPeptide, Watkinsville, GA, USA) was reconstituted to 1 mg/mL concentration and then further diluted to 0.1mg/mL for the reaction. Peroxynitrite (10 mL at 2.5 mM, for a final concentration of 500 μM) was then added to 40 mL of 0.1 mg/mL a-synuclein protein while shaking at 400 rpm. Plate was shaken for 10 minutes to allow for the nitration reaction to occur and the subsequent peroxynitrite decomposition. Nitrated a-synuclein was then pooled and frozen. BSA protein was diluted in 1X PBS pH 7.2 to 50 mg/mL concentration. Peroxynitrite (10 mL at 2.5 mM, for a final concentration of 500 μM) was then added to 40 mL protein while shaking at 400 rpm. Plate was shaken for 10 minutes to allow for the nitration reaction to occur and the subsequent peroxynitrite decomposition. Nitrated BSA was then pooled and frozen.

For *in vivo* seeding experiments, recombinant human αSyn monomer (SPR-321) and human αSyn pre-formed fibrils (hPFF; SPR-322) were purchased from StressMarq (Victoria, BC, Canada) and stored at −80 °C. For each stereotaxic injection session, a 2 mg/mL hPFF aliquot was thawed at room temperature and pulse-sonicated at 20% power (1 s on / 1 s off, 10 cycles, repeated twice) using a Fisher Scientific FB120 sonicator. The suspension was gently mixed and visually inspected to confirm dispersion. Sonicated hPFFs were kept at room temperature and used on the same day. Human αSyn monomer was thawed and injected without sonication.

### AAV vectors

Recombinant AAV2/6 vectors encoding human wild-type αSyn under the human synapsin-1 promoter were used to drive neuronal expression of human αSyn (HαSyn). The expression cassette included a woodchuck hepatitis virus post-transcriptional regulatory element (WPRE) and a polyadenylation signal. Vector production, purification and titration were performed by Revitty (Germany).

### In vitro assays

#### ELISA

Binding and specificity of 6G6 were examined on the Meso Scale Discovery (MSD, Rockville, MD, USA) electrochemiluminescent ELISA platform. To test the specificity of Tyr39-nitrated αSyn antibodies, αSyn-related biotinylated 50mer peptides were synthesized with single nitration sites on each of the four tyrosine (nY39, aa14-64; nY125, aa90-140; nY133, aa90-140; and nY136 aa90-140 custom synthesis, Abclonal). The non-nitrated Y39 50mer peptide (Abclonal) and a biotinylated unmodified full length synuclein (Anaspec, Fremont, CA, USA) were also evaluated for specificity. Test antibodies, to capture analyte, were coated overnight on a MULTI-ARRAY 1-Spot SECTOR plate (MSD) and blocked the next day in 1X casein (Vector Labs, Newark, CA, USA) for 1hr at room temperature (RT) on shaker at 750rpm. The plates were washed and incubated with peptides or protein in a titration (1 h RT, shaking at 750rpm). Plates were washed and binding detected with streptavidin sulfotag (MSD). After washing, read buffer B (MSD) was added and the plates were read with MSD Plate Reader, MESO QuickPlex SQ 120. A direct binding assay was performed using the MULTI-ARRAY 1-Spot SECTOR plates coated with a titration of chemically nitrated synuclein, chemically nitrated BSA as well as a full-length synuclein proteinnitrated at position Y39 (Mehl lab, Unnatural Protein Facility, Oregon State University, Corvallis, OR, USA). 6G6 or a general nitration antibody (3-NT, 06-285, Millipore-Sigma) was added and bound antibody detected with an anti-species sulfotag followed by addition of read buffer B and read on the MSD plate reader.

#### Western blotting

Recombinant unmodified or chemically nitrated at tyrosine 39 full-length human or mouse αSyn (Mehl lab) were resolved in a BOLT 4-12% bis-tris gradient gel (Invitrogen, USA) under reducing conditions and transferred to nitrocellulose using iBlot2 dry transfer device (Invitrogen). Blots were blocked with intercept blocking buffer (Licor, Lincoln, NB, USA) and probed with 6G6, total synuclein (BD Biosciences, #610787) or a 3-NT antibody (Millipore Sigma, 06-284) and detected with Licor secondary antibodies. Blots were imaged using an Odyssey CLX instrument (Licor).

#### Label-free binding kinetics with biolayer interferometry (BLI)

Binding affinity (K_D_) was determined based on biolayer interferometry (BLI) assays. 3µg/ml of the biotinylated anti-Rabbit Fc (Jackson Immuno Research, Cat#111-066-046) was loaded onto high precision streptavidin (SAX) Biosensors (Sartorius, Cat. 18-5117). The loaded sensors were then blocked with 100 µM dPEG-biotin acid for 10 minutes. Then the loaded sensors were used to capture test antibodies as ligands on the sensors. The antibody loaded sensors were dipped into a serial dilution of analyte (50-mer peptide nY39) at various concentrations. Reference sample well (buffer, PBS with 0.1% BSA, 0.02% Tween-20, pH 6.0) was used as control for data analysis. Kinetic constants were calculated using a monovalent (1:1) binding model whenever feasible based on the response. Binding affinity (K_D_) was calculated using the ratio of dissociation rate (k_d_)/dissociation rate (k_a_). Similar experiments were performed for mouse antibody testing but with anti-mouse capture (AMC) biosensors (Sartorius, Cat#18-5088).

#### SIMOA assays for Tyr39-nitrated and total α-synuclein

A single-molecule array (SIMOA) assay was developed on the Quanterix SR-X platform (Billerica, MA, USA) to quantify Tyr39-nitrated αSyn in human CSF. 8C2 was conjugated to SIMOA 488-dyed beads using MES pH 5.5 coupling buffer. The total αSyn antibody (BD Biosciences, #610787) was biotinylated and used as detector. For each run, 500,000 8C2-coated beads in 25 µL were incubated with 100 µL of sample or calibrator for 1 h at 30 °C (800 rpm). A synthetically prepared 70-mer αSyn peptide nitrated at Tyr39 (aa31–100; GKTKEGVL(nitrated-Y)VGSKTKEGVVHGVATVAEKTKEQVTNVGGAVVTGVTAVAQKTVEGAGSIAAAT GFVKKDQL) was used as calibrator. Human CSF samples were obtained commercially BioVT (formerly PrecisionMed). The cohort included CSF specimens from 10 individuals with a clinical diagnosis of Parkinson’s disease and 5 individuals without a Parkinson’s disease diagnosis. The non-PD samples were selected as age- and sex-matched controls where possible. Demographic and clinical characteristics, including age at baseline visit, sex, ethnicity, Mini-Mental State Examination score, Parkinson’s disease diagnosis and modified Hoehn and Yahr stage, are provided in Table 2. Samples were diluted 2–4-fold in Quanterix sample buffer. After incubation and washing, 100 µL of biotinylated detector antibody (0.6 µg/mL) was added for 20 min at 30 °C, followed by washing and 10-min incubation with 100 µL streptavidin–β-galactosidase (200 pM). After a final wash, resorufin-β-D-galactosidase substrate was added and signals were recorded as average enzymes per bead (AEB). Calibrator curves were fitted with a four-parameter logistic (4-PL) model (Quanterix software or GraphPad Prism), accepting regressions with R² > 0.95 and calibrators within 25% of nominal concentration. Unknown concentrations were interpolated and corrected for sample dilution. A total αSyn SIMOA assay was also established using bead-conjugated MJFR1 (Abcam, #ab138501) as capture antibody and the biotinylated BD antibody as detector. Total protein levels in CSF were measured with the Quant-iT assay (Invitrogen).

## Methods

### Animals

All animal experiments were conducted in accordance with national and institutional regulations. Procedures performed in Germany were approved by the Landesamt für Verbraucherschutz und Ernährung NRW (LAVE) and complied with the German Animal Welfare Act and the EU Directive 2010/63/EU. Experiments in the United States were conducted at the Nitrase Therapeutics Vivarium operated by Mispro Biotech Services (South San Francisco, CA, USA) under protocols approved by the Mispro Institutional Animal Care and Use Committee (IACUC) and in accordance with the US Animal Welfare Act and the Guide for the Care and Use of Laboratory Animals.

Female C57BL/6JRj mice (Janvier Labs), 15–22 weeks of age at surgery, were used for experiments combining vagal AAV injections and paraquat treatment. For studies modelling GBA1-associated synucleinopathy, heterozygous knock-in Gba1L444P/wt mice (B6;129S4-Gbatm1Rlp/Mmnc) and wild-type littermates were obtained from the Mutant Mouse Regional Resource Centre (MMRRC). Genotyping was performed by PCR of genomic DNA using forward primer 5′-CCCCAGATGACTGATGCTGGA-3′ and reverse primer 5′-CCAGGTCAGGATCACTGATGG-3′ followed by NciI digestion, which yielded 386 and 200 bp fragments in heterozygous animals; wild-type mice showed an undigested band. Aged (22– 30-month-old) Gba1L444P/wt and wild-type mice were used for AAV2/6-HαSyn injections.

Homozygous M83 transgenic mice (B6.Cg-Tg(Prnp-SNCA*A53T)83Vle/J; Jackson Laboratory #004479) expressing human A53T αSyn under the prion promoter were bred in-house at Nitrase Therapeutics. Male and female animals aged 8–9 weeks at surgery were used for PFF-seeding experiments. All mice were housed in individually ventilated cages in specific-pathogen-free facilities on a 12 h light/dark cycle with ad libitum access to food and water.

### Surgeries and treatments

Animals were randomly allocated to surgery and antibody treatment groups. Where possible, animals from different cages/litters were distributed across treatment groups to minimise cage and genotype related confounding. Investigators assigning animals and administering antibodies were aware of group allocation, outcome assessment and quantitative analyses were performed blinded to treatment group. For vagal AAV injections, mice were anaesthetised with isoflurane and given buprenorphine analgesia. A midline cervical incision was made, the left vagus nerve was exposed and 800 nL of AAV2/6-HαSyn was injected at 350 nL/min through a 35-gauge blunt steel needle attached to a 10 µL NanoFil syringe. The needle was left in place for 2–3 min before withdrawal to minimise reflux, and the incision was closed.

In the paraquat vagal paradigm, paraquat dichloride hydrate (Sigma-Aldrich) was dissolved in 0.9% saline and administered intraperitoneally at 5 mg/kg on days 15 and 22 after AAV injection; control mice received volume-matched saline. In both paraquat and Gba1L444P/wt experiments, treatment with 6G6 or isotype control IgG was initiated 5 days after AAV surgery. Mice received weekly intraperitoneal injections of 6G6 (100 mg/kg; 100 mg/mL formulation) or isogenic mouse IgG1 control for a total of four doses.

For PFF-seeding experiments, M83 mice were anaesthetised with isoflurane and placed in a stereotaxic frame. Animals received Meloxicam-SR (4 mg/kg, s.c.) before surgery for perioperative analgesia. A burr hole (<0.7 mm) was drilled above the right dorsal striatum and 2 µL of human αSyn PFFs or monomer (4 µg; 2 µg/µL) were injected at +1.0 mm anteroposterior, −1.5 mm mediolateral and −2.5 mm dorsoventral relative to bregma using a 33-gauge stainless-steel cannula at 300 nL/min. The cannula was left in place for 3 min before withdrawal. Antibody treatment (6G6 or mouse IgG1, 100 mg/kg, intraperitoneal) was started 2 h before PFF or monomer injection and continued once weekly for 7 weeks.

On the day of sacrifice, mice were deeply anaesthetised with sodium pentobarbital (i.p.). Immediately before perfusion, terminal blood samples were obtained by submandibular venipuncture (vagal experiments) or cardiac puncture (PFF experiments) into EDTA-coated Microvette tubes (Sarstedt). Samples were centrifuged at 5,000–10,000 g for 10 min at 4 °C, and plasma aliquots were stored at −80 °C for pharmacokinetic analyses.

Following blood collection, mice were perfused transcardially or through the ascending aorta with PBS (PFF experiments) or PBS followed by 4% (w/v) paraformaldehyde in phosphate buffer (vagal experiments). In PFF-injected M83 mice, olfactory bulbs were dissected, snap-frozen and stored at −80 °C; the remaining brain was immersion-fixed in 4% paraformaldehyde for 48 h and transferred to PBS containing 0.02% sodium azide. In vagal experiments, following 4% paraformaldehyde perfusions brains were immersed in 4% paraformaldehyde for 24 h and cryoprotected in 30% sucrose.

### Tissue processing and immunohistochemistry

For vagal AAV experiments, coronal brain sections (35 µm) were cut on a freezing microtome and collected free-floating. For brightfield immunohistochemistry, sections were quenched in 3% H_2_O_2_ and 10% methanol in TBS, blocked with 5% normal serum and incubated overnight at room temperature with rabbit anti-human αSyn (MJFR1, Abcam; 1:50,000). After TBS washes, biotinylated secondary antibodies (1:200; Vector Laboratories) and avidin–biotin– peroxidase complex (ABC Elite, Vector Laboratories) were applied sequentially. Immunoreactivity was visualised with 3,3′-diaminobenzidine (DAB) and sections were mounted and coverslipped with Depex (Sigma-Aldrich). For fluorescent detection, sections were treated with TrueBlack to reduce autofluorescence, blocked in 5% normal goat serum and incubated with anti-human αSyn (MJFR1, Abcam; 1:3000) overnight at room temperature, followed by biotinylated secondary antibody and streptavidin–DyLight 649 (Vector Laboratories). Nuclei were counterstained with DAPI and sections were coverslipped with ProLong Gold Antifade (Invitrogen).

Brains from PFF- and monomer-injected M83 mice were processed at NeuroScience Associates (Knoxville, TN, USA) using MultiBrain technology. After equilibration in 20% glycerol and 2% dimethylsulfoxide, brains were embedded in gelatin, cured with formaldehyde, frozen and coronally sectioned at 30 µm. Serial sections through the whole brain were collected. Every sixth section (180 µm interval) was processed free-floating for pSer129-αSyn and TH immunohistochemistry. After quenching and TBS rinses, sections were incubated overnight at room temperature with primary antibodies against pSer129-αSyn (Abcam, ab51253, 1:400,000) or TH (Pelfreez, P40101, 1:6000). Biotinylated secondary antibodies (Vector Laboratories) and DAB (with or without nickel enhancement) were used for visualisation. For Nissl staining, TH-immunostained sections were incubated in thionine solution (acetate buffer, pH 4.5), differentiated, dehydrated in graded ethanols, cleared in xylene and coverslipped with Permount mounting medium.

### Antibody concentration measurements in plasma and brain

6G6 concentrations in plasma (all *in vivo* studies) and olfactory bulb homogenates (PFF study) were measured using an MSD electrochemiluminescent assay. Pre-blocked MSD GOLD Streptavidin plates were coated with 30 µL of 0.5 µg/mL biotinylated nY39 50-mer peptide (aa14–64; Biotin-GVVAAAEKTKQGVAEAAGKTKEGVL(nitrated-Y)VGSKTKEGVVHGVATVAEKTKEQVT) in PBS/0.1× casein for 1 h at room temperature (750 rpm). After washing, diluted plasma (1:40,000–1:160,000) or olfactory bulb homogenates (1:10–1:40) were added together with a 6G6 standard curve and incubated for 1 h. After washing, an anti-mouse sulfotag-conjugated secondary antibody (MSD) was added for 1 h, followed by MSD Read Buffer B and reading on a MESO QuickPlex SQ 120 instrument. Concentrations were interpolated from 4-PL standard curves (GraphPad Prism) and corrected for sample dilution.

### Image analysis

HαSyn optic density was assessed in 3 equally spaced dMO sections containing DMnX and NTS. Samples were imaged using Zeiss AXIO Observer microscope at 20X magnification. The region of interest encompassing both DMnX and NTS was delineated with Fiji (ImageJ version 2.1.0/1.53). Background was subtracted using a rolling-ball algorithm (radius of 25 pixels, light background) and Gaussian blur filter (sigma = 0.5) was used.

For PFF-induced pathology, pSer129-αSyn-stained sections were scanned at 20× magnification on an Olympus VS200 slide scanner (0.274 µm/pixel). Whole-slide images were uploaded to Concentriq (Proscia) and analysed using Aiforia, a cloud-based image analysis platform. For each region (substantia nigra, thalamus), a convolutional neural-network algorithm was trained to recognise total tissue area and pSer129-positive inclusions and neurites. Training consisted of iterative cycles of manual annotation of tissue and lesion classes, algorithm training with increasing iterations and review/refinement of annotations until satisfactory performance was achieved. The final algorithms were applied bilaterally to three anatomically matched levels of the substantia nigra (SN) and thalamus per animal, with regions of interest manually traced for each section. For each animal and region, the number of pSer129-positive objects and the analysed tissue area were exported and organised in Microsoft Excel. Pathology burden was expressed as the number of pSer129-positive objects per unit area.

### Spreading analysis

For quantification of axonal spreading, the total number of HαSyn-immunoreactive axons was counted in coronal sections of the left pons at Bregma −5.40 mm. Axons were identified as winding threads with densely labelled, irregularly spaced varicosities. For quantitative measurements of axonal length and density in the ventral pons, stereological methods using the SpaceBalls probe were applied as detailed below (see Stereology).

### Stereology

Stereological analyses were used both for quantifying HαSyn-positive axonal length and density in the pons and for estimating nigral neuron numbers in the PFF model. For axonal length and density measurements, three equally spaced coronal pontine sections (Bregma−5.68, −5.51 and −5.34 mm) were analysed. A region of interest encompassing ventral pontine regions was delineated using established anatomical landmarks. The SpaceBalls stereological probe (Stereo Investigator software, MBF Bioscience) was used to obtain unbiased estimates of total HαSyn-positive axonal length and length density within the defined region. Analyses were performed on an Observer.Z1 microscope (Zeiss) equipped with a motorised stage and 63× Plan-Apochromat objective. Unbiased stereological estimation of nigral neuron numbers was performed using the optical fractionator method (Stereo Investigator 2019, MBF Bioscience). The substantia nigra pars compacta (SNc) ipsilateral to the injection site was delineated on every sixth midbrain section between Bregma −2.7 and −3.6 mm (Franklin and Paxinos, 2008), based on large, densely packed TH-immunoreactive neurons. TH-positive neurons in substantia nigra pars reticulata, substantia nigra pars lateralis and ventral tegmental area were excluded. Counting was performed at high magnification (100× UPlanSApo objective) using a 1 µm guard zone at the top and bottom of each section. The coefficient of error was calculated according to Gundersen and Jensen (1987) and was <0.10 for all estimates.

### Statistical analysis

Statistical analyses were performed using GraphPad Prism (version 10.6.0) and sample sizes were calculated using G*Power (v3.1.9.6). Data are presented as mean ± standard deviation (SD) unless otherwise indicated. For comparisons between two independent groups, unpaired two-tailed Student’s t-tests were used; when variances were unequal, Welch’s correction was applied. For comparisons involving more than two groups with a single factor, one-way ANOVA followed by Tukey’s multiple-comparisons test was used. For grouped comparisons with two, two-way ANOVA was employed, followed by Tukey’s post hoc test to compare selected group means. Statistical significance was set at p < 0.05.

## Data availability

The raw data underlying this article will be shared on reasonable request to the corresponding author.

## Acknowledgements

We thank Dr. Rita Pinto-Costa, Ms. Laura Demmer, Ms. Zoe Fisk personnel at the DZNE Light Microscope Facility, Image and Data Analysis Facility and Preclinical Center for assistance with the experiments and assays. We would also like to thank Prasanna Kandel and Frank Kayser for review of this manuscript.

## References

1. Spillantini MG, Schmidt ML, Lee VM, Trojanowski JQ, Jakes R, Goedert M. Alpha-synuclein in Lewy bodies. Nature. Aug 28 1997;388(6645):839–40. doi:10.1038/42166

2. Spillantini MG, Crowther RA, Jakes R, Hasegawa M, Goedert M. alpha-Synuclein in filamentous inclusions of Lewy bodies from Parkinson’s disease and dementia with lewy bodies. Proc Natl Acad Sci U S A. May 26 1998;95(11):6469–73. doi:10.1073/pnas.95.11.6469

3. Goedert M, Jakes R, Spillantini MG. The Synucleinopathies: Twenty Years On. J Parkinsons Dis. 2017;7(s1):S51–S69. doi:10.3233/JPD-179005

4. Braak H, Del Tredici K, Rub U, de Vos RA, Jansen Steur EN, Braak E. Staging of brain pathology related to sporadic Parkinson’s disease. Neurobiol Aging. Mar-Apr 2003;24(2):197–211. doi:10.1016/s0197-4580(02)00065-9

5. Del Tredici K, Braak H. Review: Sporadic Parkinson’s disease: development and distribution of alpha-synuclein pathology. Neuropathol Appl Neurobiol. Feb 2016;42(1):33–50. doi:10.1111/nan.12298

6. Brundin P, Melki R, Kopito R. Prion-like transmission of protein aggregates in neurodegenerative diseases. Nat Rev Mol Cell Biol. Apr 2010;11(4):301–7. doi:10.1038/nrm2873

7. Luk KC, Kehm V, Carroll J, et al. Pathological alpha-synuclein transmission initiates Parkinson-like neurodegeneration in nontransgenic mice. Science. Nov 16 2012;338(6109):949–53. doi:10.1126/science.1227157

8. Peelaerts W, Bousset L, Van der Perren A, et al. alpha-Synuclein strains cause distinct synucleinopathies after local and systemic administration. Nature. Jun 18 2015;522(7556):340–4. doi:10.1038/nature14547

9. Sacino AN, Brooks M, Thomas MA, et al. Intramuscular injection of alpha-synuclein induces CNS alpha-synuclein pathology and a rapid-onset motor phenotype in transgenic mice. Proc Natl Acad Sci U S A. Jul 22 2014;111(29):10732–7. doi:10.1073/pnas.1321785111

10. Prusiner SB, Woerman AL, Mordes DA, et al. Evidence for alpha-synuclein prions causing multiple system atrophy in humans with parkinsonism. Proc Natl Acad Sci U S A. Sep 22 2015;112(38):E5308–17. doi:10.1073/pnas.1514475112

11. Lloyd GM, Sorrentino ZA, Quintin S, et al. Unique seeding profiles and prion-like propagation of synucleinopathies are highly dependent on the host in human alpha-synuclein transgenic mice. Acta Neuropathol. Jun 2022;143(6):663–685. doi:10.1007/s00401-022-02425-4

12. Luk KC, Kehm VM, Zhang B, O’Brien P, Trojanowski JQ, Lee VM. Intracerebral inoculation of pathological alpha-synuclein initiates a rapidly progressive neurodegenerative alpha-synucleinopathy in mice. J Exp Med. May 7 2012;209(5):975–86. doi:10.1084/jem.20112457

13. Ulusoy A, Rusconi R, Perez-Revuelta BI, et al. Caudo-rostral brain spreading of alpha-synuclein through vagal connections. EMBO Mol Med. Jul 2013;5(7):1119–27. doi:10.1002/emmm.201302475

14. Helwig M, Klinkenberg M, Rusconi R, et al. Brain propagation of transduced alpha-synuclein involves non-fibrillar protein species and is enhanced in alpha-synuclein null mice. Brain. Mar 2016;139(Pt 3):856–70. doi:10.1093/brain/awv376

15. Kalia M, Sullivan JM. Brainstem projections of sensory and motor components of the vagus nerve in the rat. J Comp Neurol. Nov 1 1982;211(3):248–65. doi:10.1002/cne.902110304

16. Pinto-Costa R, Harbachova E, La Vitola P, Di Monte DA. Overexpression-Induced alpha-Synuclein Brain Spreading. Neurotherapeutics. Jan 2023;20(1):83–96. doi:10.1007/s13311-022-01332-6

17. La Vitola P, Szego EM, Pinto-Costa R, et al. Mitochondrial oxidant stress promotes alpha-synuclein aggregation and spreading in mice with mutated glucocerebrosidase. NPJ Parkinsons Dis. Dec 11 2024;10(1):233. doi:10.1038/s41531-024-00842-8

18. Giasson BI, Duda JE, Murray IV, et al. Oxidative damage linked to neurodegeneration by selective alpha-synuclein nitration in synucleinopathy lesions. Science. Nov 3 2000;290(5493):985–9. doi:10.1126/science.290.5493.985

19. Betarbet R, Sherer TB, MacKenzie G, Garcia-Osuna M, Panov AV, Greenamyre JT. Chronic systemic pesticide exposure reproduces features of Parkinson’s disease. Nat Neurosci. Dec 2000;3(12):1301–6. doi:10.1038/81834

20. Tieu K, Perier C, Caspersen C, et al. D-beta-hydroxybutyrate rescues mitochondrial respiration and mitigates features of Parkinson disease. J Clin Invest. Sep 2003;112(6):892–901. doi:10.1172/JCI18797

21. Dias V, Junn E, Mouradian MM. The role of oxidative stress in Parkinson’s disease. J Parkinsons Dis. 2013;3(4):461–91. doi:10.3233/JPD-130230

22. Danielson SR, Held JM, Schilling B, Oo M, Gibson BW, Andersen JK. Preferentially increased nitration of alpha-synuclein at tyrosine-39 in a cellular oxidative model of Parkinson’s disease. Anal Chem. Sep 15 2009;81(18):7823–8. doi:10.1021/ac901176t

23. Hodara R, Norris EH, Giasson BI, et al. Functional consequences of alpha-synuclein tyrosine nitration: diminished binding to lipid vesicles and increased fibril formation. J Biol Chem. Nov 12 2004;279(46):47746–53. doi:10.1074/jbc.M408906200

24. Paxinou E, Chen Q, Weisse M, et al. Induction of alpha-synuclein aggregation by intracellular nitrative insult. J Neurosci. Oct 15 2001;21(20):8053–61. doi:10.1523/JNEUROSCI.21-20-08053.2001

25. Yu Z, Xu X, Xiang Z, et al. Nitrated alpha-synuclein induces the loss of dopaminergic neurons in the substantia nigra of rats. PLoS One. Apr 8 2010;5(4):e9956. doi:10.1371/journal.pone.0009956

26. Musgrove RE, Helwig M, Bae EJ, et al. Oxidative stress in vagal neurons promotes parkinsonian pathology and intercellular alpha-synuclein transfer. J Clin Invest. Jun 13 2019;129(9):3738–3753. doi:10.1172/JCI127330

27. Helwig M, Ulusoy A, Rollar A, et al. Neuronal hyperactivity-induced oxidant stress promotes in vivo alpha-synuclein brain spreading. Sci Adv. Sep 2 2022;8(35):eabn0356. doi:10.1126/sciadv.abn0356

28. Wright S, Dang VC, Hussain S, et al. Selective peroxynitrite-mediated protein nitration catalyzed by glyoxalase domain containing protein 4. Proc Natl Acad Sci U S A. Feb 10 2026;123(6):e2515002123. doi:10.1073/pnas.2515002123

29. Sarah Wright VCD, Sami Hussain, Prasanna Kandel, Robert P. Brendza, Sahar Mazhar Marie Whitmore, Selim Boudoukha, Jaskamal Banwait, Edward Vertudes, Kate Markham, Marta Trzeciak, Grace Pohan, Andy Jennings, Sheerin Shahidi-Latham, Frank Kayser, Mike Beckstead, Aaron L. Lucius, Arun Kashyap, Harry Ischiropoulos, Irene Griswold-Prenner. A novel function of glyoxalase domain containing protein 4 (GLOD4) is associated with1neuron dysfunction and neurodegeneration2. Research Square. 2025;10.21203/rs.3.rs-6100422/v1

30. Burai R, Ait-Bouziad N, Chiki A, Lashuel HA. Elucidating the Role of Site-Specific Nitration of alpha-Synuclein in the Pathogenesis of Parkinson’s Disease via Protein Semisynthesis and Mutagenesis. J Am Chem Soc. Apr 22 2015;137(15):5041–52. doi:10.1021/ja5131726

31. Porter JJ, Jang HS, Van Fossen EM, et al. Genetically Encoded Protein Tyrosine Nitration in Mammalian Cells. ACS Chem Biol. Jun 21 2019;14(6):1328–1336. doi:10.1021/acschembio.9b00371

32. Wu S, Huang HW, Panchal A, Chowdhury EA, Shah DK. Quantitation of regional distribution of antibodies in rat brain following systemic and intra-CNS administration. J Cereb Blood Flow Metab. Sep 2025;45(9):1785–1798. doi:10.1177/0271678X251333536

33. Tolve M, Ulusoy A, Patikas N, et al. The transcription factor BCL11A defines distinct subsets of midbrain dopaminergic neurons. Cell Rep. Sep 14 2021;36(11):109697. doi:10.1016/j.celrep.2021.109697

34. Volpicelli-Daley LA, Luk KC, Patel TP, et al. Exogenous alpha-synuclein fibrils induce Lewy body pathology leading to synaptic dysfunction and neuron death. Neuron. Oct 6 2011;72(1):57–71. doi:10.1016/j.neuron.2011.08.033

35. Paumier KL, Luk KC, Manfredsson FP, et al. Intrastriatal injection of pre-formed mouse alpha-synuclein fibrils into rats triggers alpha-synuclein pathology and bilateral nigrostriatal degeneration. Neurobiol Dis. Oct 2015;82:185–199. doi:10.1016/j.nbd.2015.06.003

36. Schildknecht S, Gerding HR, Karreman C, et al. Oxidative and nitrative alpha-synuclein modifications and proteostatic stress: implications for disease mechanisms and interventions in synucleinopathies. J Neurochem. May 2013;125(4):491–511. doi:10.1111/jnc.12226

37. Di Monte DA. The environment and Parkinson’s disease: is the nigrostriatal system preferentially targeted by neurotoxins? Lancet Neurol. Sep 2003;2(9):531–8. doi:10.1016/s1474-4422(03)00501-5

38. Grassi D, Howard S, Zhou M, et al. Identification of a highly neurotoxic alpha-synuclein species inducing mitochondrial damage and mitophagy in Parkinson’s disease. Proc Natl Acad Sci U S A. Mar 13 2018;115(11):E2634–E2643. doi:10.1073/pnas.1713849115

39. Geibl FF, Musa AAS, Dietrich L, et al. Mapping cellular vulnerability in Parkinson’s disease using retro-AAVs and preformed alpha-synuclein fibrils. Transl Neurodegener. Jan 30 2026;15(1):2. doi:10.1186/s40035-026-00535-7

40. Dauer W, Przedborski S. Parkinson’s disease: mechanisms and models. Neuron. Sep 11 2003;39(6):889–909. doi:10.1016/s0896-6273(03)00568-3

41. Xie YX, Naseri NN, Fels J, et al. Lysosomal exocytosis releases pathogenic alpha-synuclein species from neurons in synucleinopathy models. Nat Commun. Aug 22 2022;13(1):4918. doi:10.1038/s41467-022-32625-1

42. Scheiblich H, Eikens F, Wischhof L, et al. Microglia rescue neurons from aggregate-induced neuronal dysfunction and death through tunneling nanotubes. Neuron. Sep 25 2024;112(18):3106–3125 e8. doi:10.1016/j.neuron.2024.06.029

43. Prymaczok NC, De Francesco PN, Mazzetti S, et al. Cell-to-cell transmitted alpha-synuclein recapitulates experimental Parkinson’s disease. NPJ Parkinsons Dis. Jan 6 2024;10(1):10. doi:10.1038/s41531-023-00618-6

44. Wu S, Hernandez Villegas NC, Sirkis DW, Thomas-Wright I, Wade-Martins R, Schekman R. Unconventional secretion of alpha-synuclein mediated by palmitoylated DNAJC5 oligomers. Elife. Jan 10 2023;12doi:10.7554/eLife.85837

45. Lang AE, Siderowf AD, Macklin EA, et al. Trial of Cinpanemab in Early Parkinson’s Disease. N Engl J Med. Aug 4 2022;387(5):408–420. doi:10.1056/NEJMoa2203395

46. Pagano G, Taylor KI, Anzures-Cabrera J, et al. Trial of Prasinezumab in Early-Stage Parkinson’s Disease. N Engl J Med. Aug 4 2022;387(5):421–432. doi:10.1056/NEJMoa2202867

47. Pagano G, Monnet A, Reyes A, et al. Sustained effect of prasinezumab on Parkinson’s disease motor progression in the open-label extension of the PASADENA trial. Nat Med. Dec 2024;30(12):3669–3675. doi:10.1038/s41591-024-03270-6

48. Buur L, Wiedemann J, Larsen F, et al. Randomized Phase I Trial of the alpha-Synuclein Antibody Lu AF82422. Mov Disord. Jun 2024;39(6):936–944. doi:10.1002/mds.29784

49. Kallunki P, Sotty F, Willen K, et al. Rational selection of the monoclonal alpha-synuclein antibody amlenetug (Lu AF82422) for the treatment of alpha-synucleinopathies. NPJ Parkinsons Dis. May 22 2025;11(1):132. doi:10.1038/s41531-024-00849-1

50. Shering C, Pomfret M, Kubiak RJ, et al. Phase 1, randomized trials of MEDI1341: cerebrospinal fluid free alpha-synuclein lowered by >50. Brain Commun. 2025;7(5):fcaf304. doi:10.1093/braincomms/fcaf304

51. Zachrisson O, Johannesson M, Soderberg L, et al. Exidavnemab binds to aggregated alpha-synuclein in human brains affected by alpha-synucleinopathies. Neurotherapeutics. Jan 2026;23(1):e00779. doi:10.1016/j.neurot.2025.e00779

52. Weihofen A, Liu Y, Arndt JW, et al. Development of an aggregate-selective, human-derived alpha-synuclein antibody BIIB054 that ameliorates disease phenotypes in Parkinson’s disease models. Neurobiol Dis. Apr 2019;124:276–288. doi:10.1016/j.nbd.2018.10.016

53. Games D, Valera E, Spencer B, et al. Reducing C-terminal-truncated alpha-synuclein by immunotherapy attenuates neurodegeneration and propagation in Parkinson’s disease-like models. J Neurosci. Jul 9 2014;34(28):9441–54. doi:10.1523/JNEUROSCI.5314-13.2014

54. Xiao B, Tan EK. Prasinezumab slows motor progression in Parkinsons disease: beyond the clinical data. NPJ Parkinsons Dis. Feb 19 2025;11(1):31. doi:10.1038/s41531-025-00886-4

55. Rubilar JC, Outeiro TF, Klein AD. The lysosomal beta-glucocerebrosidase strikes mitochondria: implications for Parkinson’s therapeutics. Brain. Aug 1 2024;147(8):2610–2620. doi:10.1093/brain/awae070

